# Simulation-conditioned generative modeling for biologically realistic pattern prediction

**DOI:** 10.64898/2025.12.13.694038

**Authors:** Kinshuk Sahu, Harris M. Davis, Jia Lu, César A. Villalobos, Avi Heyman, Emrah Şimşek, Lingchong You

## Abstract

Pattern formation underlies biological organization across scales, but predicting experimentally observed patterns remains difficult because mechanistic models and data-driven generative models fail in complementary ways. Coarse-grained mechanistic models can encode causal constraints and global morphology, yet they omit fine-scale features such as texture, color gradients, and stochastic replicate-to-replicate variation. In contrast, generative image models can produce realistic images but are not inherently grounded in the biophysical rules that shape real patterns. Here, we introduce a simulation-conditioned generative framework that uses mechanistic simulations as spatial priors for generating biologically realistic pattern data. As a concrete test, we use a synthetic-to-real inverse task to show that these generated patterns preserve information needed for inference on real experimental images, beyond merely reproducing plausible visual appearance. Using branching colony expansion of Pseudomonas aeruginosa as a model system, we combine a coarse-grained PDE model with latent representations from a foundation image model and a conditional diffusion model. The resulting framework preserves the global structures imposed by simulation while restoring experimentally observed fine-scale morphology and stochastic variability. A model trained exclusively on simulation-conditioned synthetic patterns transfers without fine-tuning to real experimental patterns, enabling inference of initial seeding configurations from experimental colony morphology. Together, these results establish simulation-conditioned generative modeling as a strategy for converting coarse mechanistic models into scientifically structured synthetic data, enabling inference tasks on real biological patterns where experimental data are scarce.

## Introduction

Self-organized pattern formation governs the arrangement and organization of biomolecules, cells, and tissues that define the structure and function of biological organisms or systems^1–3^, spanning a wide range of time (seconds to years) and length (sub-micron to kilometer) scales^4,5^. Examples include formation of division ring in a bacterial cell^6^, animal developmental patterning^7^, bacterial colony growth^8^, branching morphogenesis in tissue development^9^, and macroscopic ecosystems^10^. These patterns are not merely visual outcomes of biological dynamics; they encode the growth, motility, communication, and environmental interactions that generate them. Predicting such patterns is therefore central both to understanding how biological systems organize themselves and to engineering spatial behaviors in synthetic biology, tissue morphogenesis, and microbial communities.

For decades, extensive efforts have been made to dissect and test the mechanisms underlying pattern formation^1–3,11^. By asking whether proposed mechanisms can reproduce observed patterns and predict responses to perturbations, models provide a way to evaluate whether current biological understanding is sufficient ^5,12–16^. This predictive role is especially important for efforts to engineer spatial organization, including programming pattern formation with synthetic gene circuits ^17–19^, assembling interacting cell populations^20–22^, or controlling environmental factors^13,19^. In each case, the goal is not simply to reproduce a pattern qualitatively, but to predict how defined biological rules and initial conditions give rise to experimentally observable spatial organization.

To date, our ability to predict self-organized pattern formation has remained limited^4^. Typically these dynamics are modeled as reaction-diffusion systems using partial differential equations (PDEs)^2^ or agent-based modeling (ABM)^23^. Both approaches rely on coarse-grained representations of underlying processes, including cell growth, motility, gene expression, and, in ABMs, physical interactions between cells. Such coarse graining is often necessary because the underlying biology is incompletely known and because fully resolved simulations would be computationally prohibitive. As a result, these models require simplifying assumptions about interactions, initial conditions, and boundary conditions ^12,24^. Moreover, numerical solutions can remain computationally intensive when used for parameter sweeps or large sets of initial conditions ^17,25–27^. Despite these limitations, mechanistic models can often capture important global or topological features of biological patterns ^4,13,21,28^, such as the extent and branching geometry of expanding bacterial colonies^15^. What they usually lack is an experimentally realistic observation layer: the fine-scale morphology, texture, color, and stochastic variability produced by microscale processes such as heterogeneous motility, pigment production, and variable surface wetness^29^. Thus, coarse simulations can provide a mechanistically meaningful spatial scaffold, but they do not by themselves reproduce the full experimental pattern.

Yet, these features often carry critical information associated with strain identity^30,31^, physiological state^32^, or environmental history^33^. Variations in growth, motility, and pigment production can produce large or subtle differences in bacterial colony patterns^34,35^, while differences in carbon and nitrogen utilization can also alter colony morphology^36^. Under the same environmental condition, strains with small genetic differences can form patterns that look similar but differ in colony size, color, shape, and texture^37,38^ or transition between distinct morphotypes ^39,40^. In multi-species communities, pattern variation can reflect community interactions, spatial genetic drift, resource metabolism, growth differences, and motility variation ^21,41–43^. Thus, experimentally observed morphology is not merely a visual endpoint; it is an information-rich readout of biological state and history. Reproducing this morphology is thus important for interpreting experimental patterns and for downstream tasks such as strain identification, perturbation inference, environmental sensing, phenotyping, and the design of synthetic spatial behaviors ^38,44^.

Scientific machine learning has increasingly combined mechanistic modeling with machine learning to improve prediction, interpretability, and data efficiency. In physics-informed and biologically informed neural networks, governing equations or biologically motivated equation forms are incorporated into neural-network training to learn solutions, parameters, or missing terms from sparse data^45–48^. Related surrogate-modeling approaches train neural networks on mechanistic simulations to accelerate parameter sweeps, inverse searches, or design-space exploration. These strategies are powerful when the goal is to infer or accelerate the underlying dynamical model ^17,25–27^. However, they do not directly address a different limitation of coarse-grained simulations: the absence of experimentally realistic morphology. A fast or well-calibrated simulator can still produce patterns that preserve only global structure while lacking the morphology, texture, color, and stochastic variability present in real experiments.

Advances in generative AI have demonstrated powerful capabilities for producing realistic images from high-level prompts or conditioning inputs^49–51^. In scientific and biomedical imaging, related approaches have enabled virtual staining^52^, image-to-image translation^53,54^, and synthetic-data generation for downstream tasks such as segmentation^55^ and phenotyping ^56,57^. These studies show that realistic generated images can be useful for biological inference, especially when experimental data are limited. However, most existing approaches either translate one real image modality into another^53^, generate images from human-defined masks or annotations^55^, or use synthetic images designed primarily to improve a specific downstream task^58^. They generally do not use mechanistic simulations as spatial priors that encode how biological patterns arise from defined initial conditions or physical rules. As a result, generated images may appear realistic, but their realism is not coupled to the mechanistic dependencies that govern the biological system. This disconnect limits their use for scientific prediction, where generated outputs must remain faithful not only to image statistics but also to the mechanisms that produce the pattern ^59–61^.

Together, these limitations motivate a hybrid strategy that maintains mechanistic interpretability while leveraging the representational power of modern foundation models. Rather than embedding biophysical rules within the neural network, we present a complementary approach: using mechanistic simulations as spatial conditioning priors for a foundation image model^51^. This simulation-conditioned approach preserves the global structures dictated by a coarse-grained PDE model while allowing a generative model with high expressive capacity to render fine-scale morphological features and stochastic variability characteristic of experimental observations. Because the mechanistic and generative components are decoupled in implementation, the framework remains modular and flexible. This design enables biologically grounded, experimentally realistic pattern generation that neither mechanistic models nor generative models can achieve alone. We further use a synthetic-to-real inverse task as a functional test of the generated patterns: a model trained only on simulation-conditioned synthetic patterns is evaluated directly on real experimental colony images. We find that simulation-conditioned synthetic patterns preserve morphology-linked information sufficient for inference on real experimental images, enabling zero-shot recovery of initial seeding configurations from colony morphology.

## Results

Our framework requires two components: a mechanistic model that captures the global structure of *P. aeruginosa* branching patterns, and a learned mapping that converts coarse simulations into synthetic data structured enough to support inference on real experimental patterns. For the first component, we adopted a previously developed coarse-grained PDE model^15^. For the second, we used the Stable Diffusion Variational Autoencoder (SD VAE^51^) to obtain compact, information-rich representations of both simulated and experimental patterns, enabling efficient downstream learning.

Our hybrid strategy is conceptually different from other approaches in scientific machine learning (**Fig 1a**). Our analysis is organized around a functional criterion: whether a coarse mechanistic simulator can be converted into a source of synthetic experimental data that supports inference on real biological patterns (**Fig 1b**). To reach this test, we first evaluate whether a foundation image model provides useful latent representations of simulated and experimental patterns, then determine the data requirements for structured image-to-image translation and finally learn a simulation-to-experiment generative mapping. We then evaluate the resulting synthetic patterns in a synthetic-to-real inverse task, asking whether a model trained only on simulation-conditioned generated patterns can infer initial seeding configurations from real experimental colony images.

**Figure 1:**
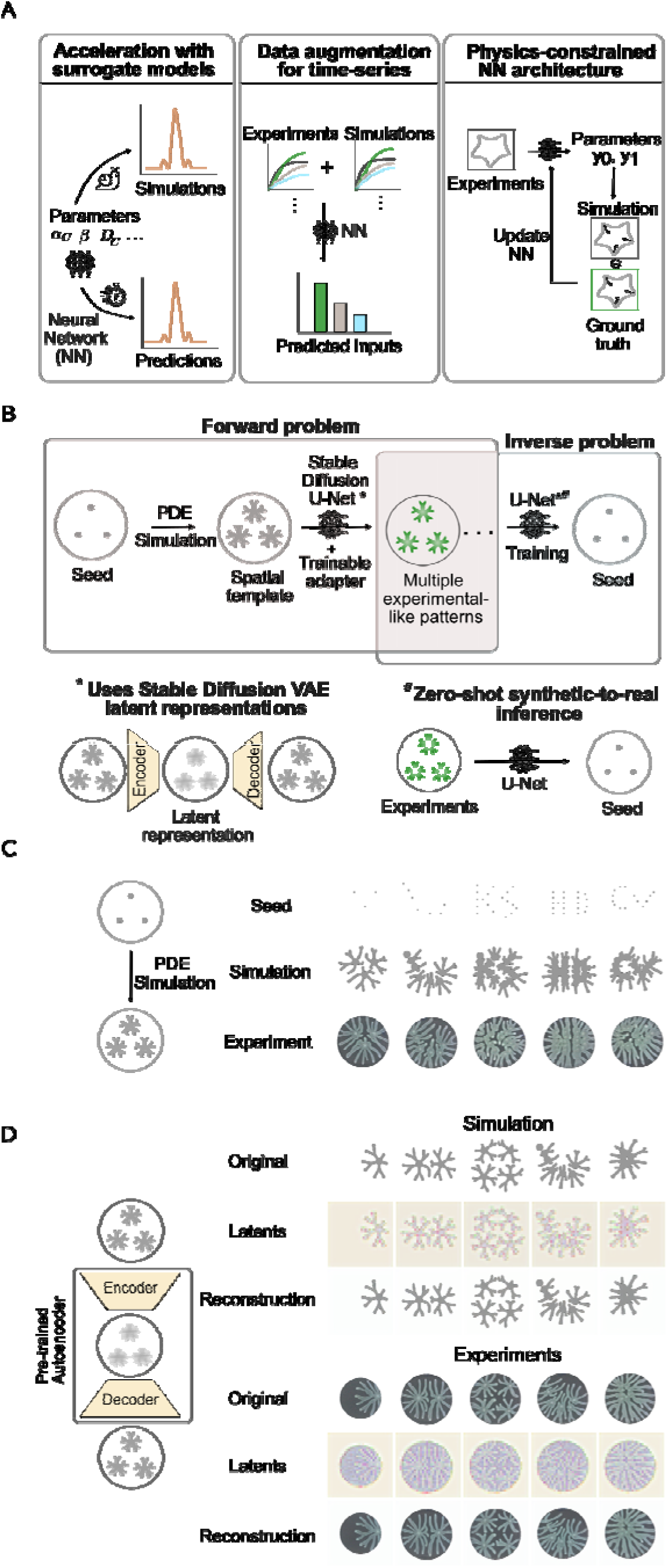
Overview of simulation-conditioned generative modelling framework and SD VAE representation on bacterial colony patterns. **(A)** *Machine learning for scientific applications: Existing* approaches commonly use machine learning to accelerate mechanistic models with surrogate neural networks, augment limited experimental datasets with simulations to train a downstream model, or include physical constraints in the neural network architecture. Recent examples include i) Building a surrogate model to rapidly screen a large parameter space for identifying desired patterning conditions^17^ (left) ii) Augmenting experimental time-series data with ODE simulations to train a ML model to infer input signals from bacterial growth and fluorescence curves^65^ (middle). iii) Employing a Physics- Bottleneck Neural Network architecture to impose hard physical constraints on inferring adhesion forces from fluorescence images in fibroblasts^46^ (right). **(B)** *Overview of our simulation-conditioned generative modelling approach:* We employed a coarse-grained PDE model adapted from Luo et al ^15^ to generate bacterial colony patterns under different initial seeding conditions(top left). These patterns serve as the spatial template for a pre-trained Stable Diffusion U-Net^51^ and trainable ControlNet adapter^73^(top middle) generative framework, operating in the SD-VAE latent space(bottom left). The framework outputs multiple experimental-like colony patterns that retain the global structure of the simulations while also capturing fine-scale morphology and underlying biological heterogeneity(top middle). We then tested the framework functionally using an inverse task, in which a predictor is trained solely on synthetic patterns to infer the initial seeds (top right) and evaluated on real experimental images without additional fine-tuning (bottom right). **(C)** *Simulation dataset captures global experimental patterning features under different initial seeding conditions*: A PDE model^15^ was used to generate *P. aeruginosa* branching patterns by varying only the initial seeding configurations. Configurations with few seeds in close proximity (e.g. 3 seeds, first column) yield long tendrils with multiple branching events, whereas high-density seeding (e.g. 20 seeds, fourth column) produces shorter but denser branches. The simulations reproduce the global morphology of experimental patterns, with local differences. **(D)** *SD VAE provides high-fidelity reconstruction of simulated and experimental patterns:* SD VAE compresses each 256×256×3 image by a factor of 48 while preserving key structural features. Top rows show the original patterns, middle rows show the latent representations, and bottom rows show the reconstructed patterns.

### A foundation image model provides compact representations of simulated and experimental bacterial patterns

Our previous study^15^ showed that a coarse-grained PDE model captures key structural characteristics of experimental branching patterns. Here, we focused on patterns emerging from varying initial spatial arrangements of seeding cells (**Figure 1c**). The complexity and diversity of these patterns make the system a suitable testbed for integrating mechanistic models with deep learning for predictive tasks.

Our experimental and simulated images are high-dimensional 256 × 256 × 3 images, motivating the use of a dimensionality-reduction tool that preserves essential information associated with these patterns for downstream tasks. The SD VAE is a pre-trained foundation image model^51^ that encodes images into latent representations and reconstructs them with high fidelity. VAEs use an encoder-bottleneck-decoder architecture (**Figure 1d**): the encoder compresses high-dimensional images into a lower-dimensional latent representation, and the decoder reconstructs the image from this bottleneck. In biological contexts, VAEs have been used to represent microbial growth curves^62–64^, fluorescence images^65^, metabolomics profiles^66^, de novo generation of antibiotics^67^, protein design^68^, and synthetic pattern formation^17^, often performing as well as or better than raw data for downstream tasks^62^. Although the SD VAE was not trained specifically on bacterial colony images, it has shown useful out-of-the-box generalization to several biological imaging domains ^57,69^. In our application, the SD VAE compressed each 256 × 256 × 3 image into a 32 × 32 × 4 latent representation, corresponding to a 48-fold reduction in dimensionality (**Figure 1d**). Reconstructions retained high fidelity, with Structural Similarity Index Measure (SSIM) scores above 0.90 (**Supplementary Figure 1**).

### Latent-space surrogates accelerate pattern predictions but inherit mechanistic simulator limitations

If the SD VAE latents preserve essential structural features of simulated patterns, they should support biologically meaningful prediction tasks. As a case study, we used the latents to accelerate mechanistic simulations via a neural network surrogate. The surrogate maps initial seeding configurations (**Supplementary Figure 2**) to the SD VAE latents (**Supplementary Figures 3 & 4**), from which the decoder reconstructs the full pattern. This approach bypasses PDE solving, enabling rapid pattern prediction.

We implemented a dilated ResNet (**Supplementary Figure 5**), inspired by surrogates in physical systems^26^, trained on 30,000 samples (90/10 training-validation split), with the SD VAE fixed (**Figure 2a, Methods**). Predictions from the ResNet-decoder combination matched ground truth (**Figure 2b**), achieving SSIM >0.7 across diverse initial conditions, including dense seeding (**Supplementary Figures 6 & 7**), indicating model generalizability. The surrogate achieved ∼1500-fold acceleration: predicting one pattern required 0.12s on CPU, compared to ∼3 minutes for a PDE simulation. Thus, latent-space surrogates can accelerate mechanistic simulations, but they do not address the missing experimental morphology that separates coarse simulations from real colony patterns. This limitation is critical because the missing morphology is not simply cosmetic; it can determine whether synthetic patterns contain the information needed for downstream inference on real experimental images.

**Figure 2:**
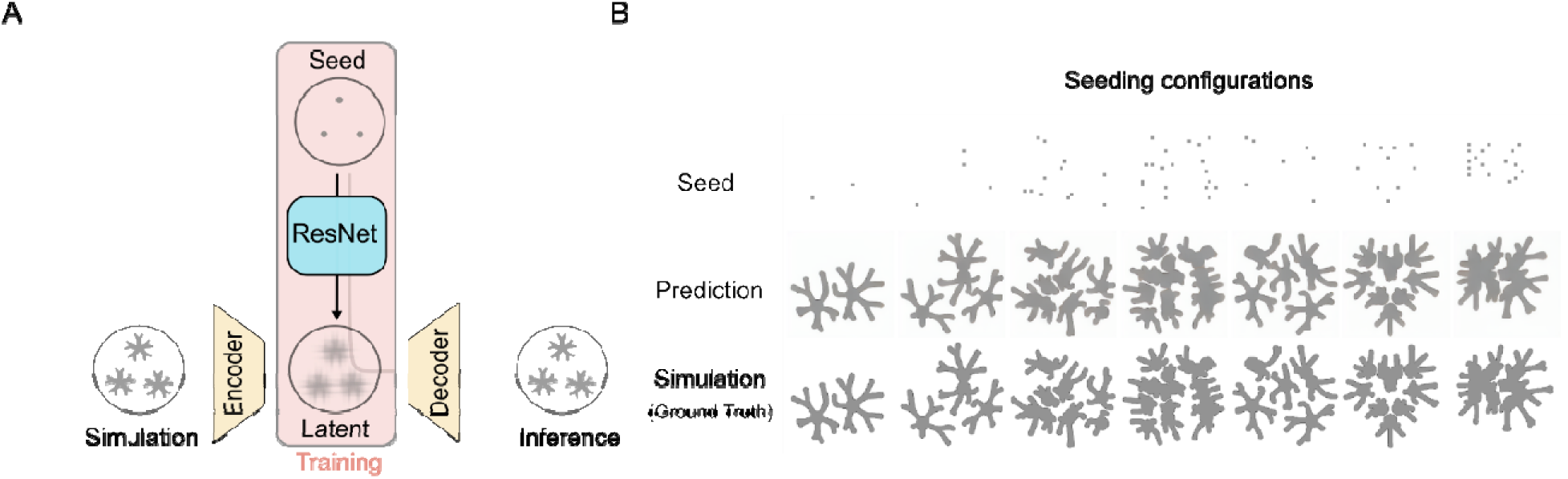
A latent-space deep-learning framework enables accelerated and reliable prediction of branching pattern formation. **(A)** *Training pipeline:* A custom dilated ResNet is trained to map initial seeding configurations to the latent representations of simulated patterns. The ResNet (outlined in blue) is trained on 30,000 seed–pattern pairs using MSE loss between predicted and ground-truth latents. The SD VAE encoder and decoder (outlined in yellow) remain fixed during training and are not updated. **(B)** *Model performance:* The trained ResNet is evaluated on an out-of-sample test set containing fixed seeding grids (top row) distinct from those used for training. Predicted latent representations are decoded into full images and compared with the corresponding PDE simulations. The surrogate model reliably reproduces the simulated patterns (compare the middle and bottom rows).

### Latent-space mapping estimates data requirements for structured image-to-image translation

Coarse-grained simulations capture the global characteristics of experimental patterns but lack detailed local features. In the simulation, branching bifurcations occur deterministically, whereas in experiments branching triggers are noisy and vary between replicates. Consequently, the PDE model generates the same pattern for each seeding configuration; but experiments generate similar yet unique patterns between replicates. This stochasticity, coupled with local morphological differences, makes direct simulation-to-experiment mapping a challenging problem.

Because experimental patterning assays are low throughput, we sought to estimate the dataset size needed to train a high-fidelity image-to-image mapping in a tractable setting. As a proxy, we examined mapping between two PDE parameter regimes that differ in global and local morphological characteristics. While this analysis does not address experimental stochasticity, it provides a controlled test for assessing the data requirements of complex image-to-image translation when both input and output patterns are high-dimensional and structurally distinct. These results serve as a lower bound on the number of paired examples likely needed for the more difficult simulation-to-experiment mapping.

We selected two PDE parameter configurations (**Methods**): the default and one that generates thinner but denser patterns (**Figure 3a**, **Supplementary Figure 8**). For a baseline, we trained on 30,000 paired simulations. Patterns from each condition were encoded into SD VAE latents (**Supplementary Figure 9**), and a ResNet was trained to map latents from one regime to the other.

**Figure 3:**
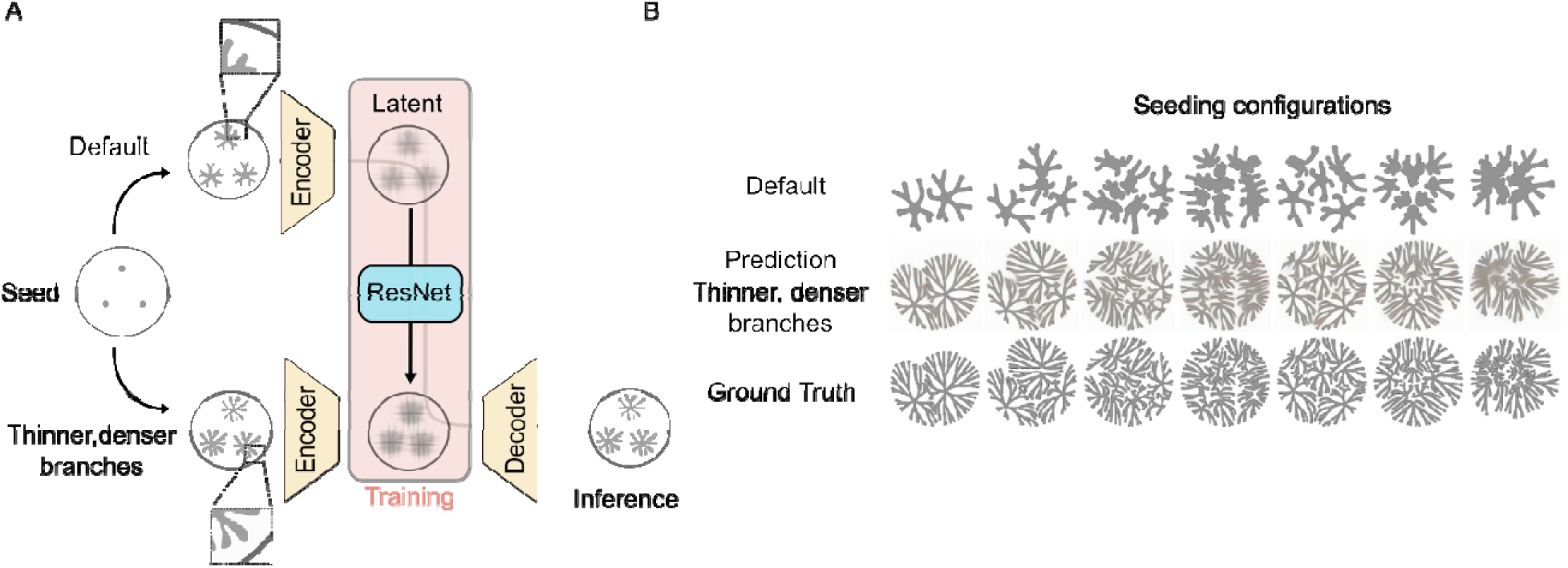
Latent-space mapping between PDE parameter regimes. **(A)** *Schematic of parameter-regime mapping.* We generated two sets of simulations using the same randomized seeding grids but different PDE parameter configurations. The second parameter set produces thinner, denser, and more intricately branched patterns. A ResNet was trained to map the latent representations from parameter set 1 to those of parameter set 2. **(B)** *Latent-space mapping captures global structural trends.* For multiple seeding configurations (columns), patterns from parameter set 1 (top row) were translated by the ResNet into predictions for parameter set 2 (middle row) and compared with the ground-truth simulations for that regime (bottom row). The model recapitulates major global features, including colony extent, branch number, and overall geometry, though some fine-scale local details near colony edges remain imperfectly captured.

The trained network achieved high accuracy on the test set (**Figure 3b**), comparable to that achieved by the seeding-to-pattern emulator, with minor differences likely reflecting the more intricate branching geometry of the second parameter regime (**Supplementary Figure 10**). We then varied the training data size between 100 and 12,800 pairs. Validation MSE decreased rapidly up to ∼1600 samples, after which performance gains diminished (**Figure 4a**). Qualitatively, high-fidelity structured patterns emerged at ∼3,200 samples, whereas models trained on ≤800 samples generated blurred outputs (**Figure 4b**). Collecting ∼3,200 experimental patterning images is feasible but experimentally laborious and time consuming. We then evaluated whether we could use data augmentation to lower the data demand. Since the patterns are rotationally invariant, we augmented the dataset to 40,000 samples by applying rotations to the paired simulation patterns. With rotational augmentation, non-blurred predictions emerged with approximately 80 unique paired patterns (Figure 4), providing a practical lower-bound estimate for the experimental scale needed to establish simulation-to-experiment mapping in this system.

**Figure 4.**
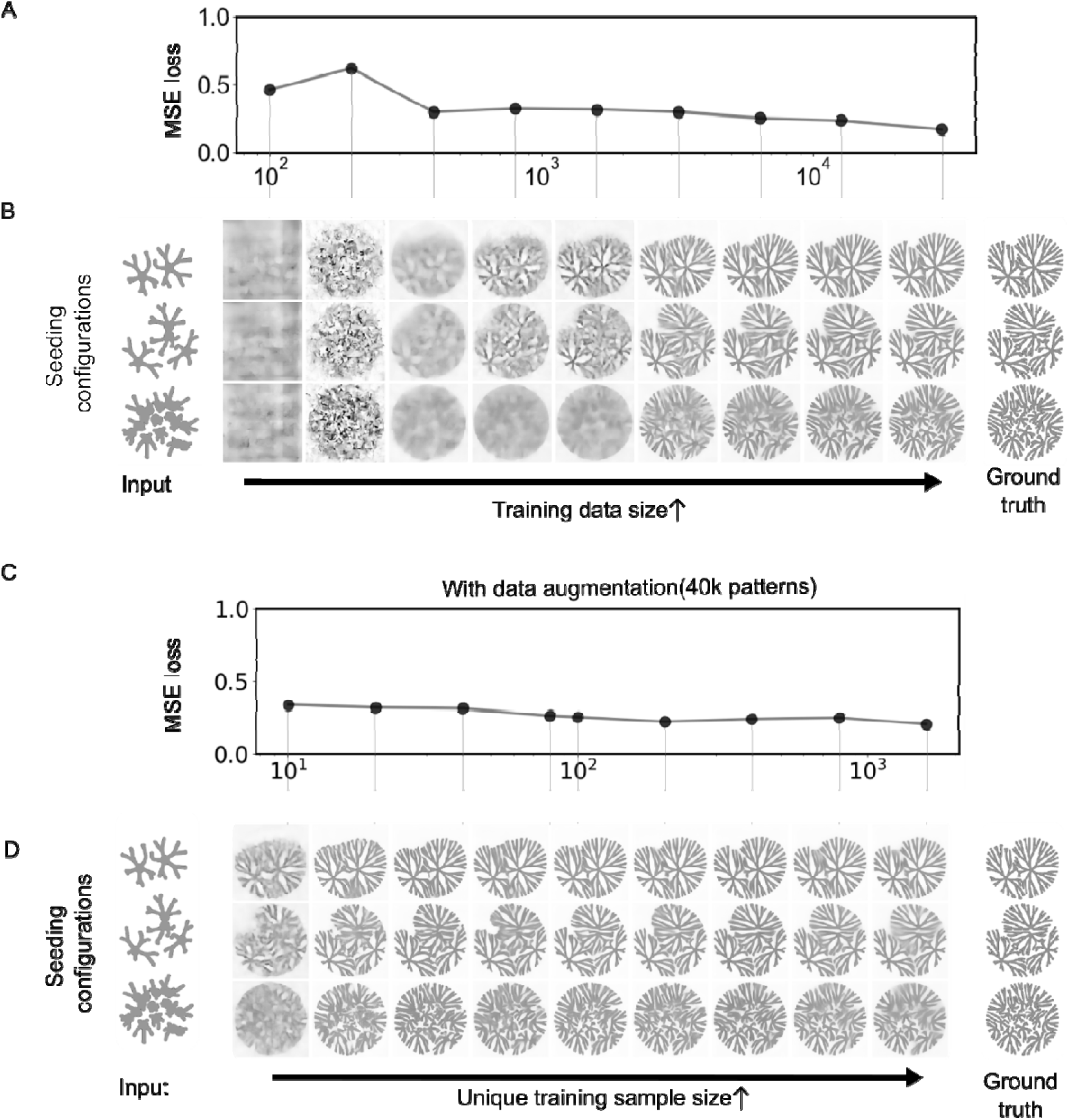
Data augmentation substantially reduces the number of unique samples required for high-fidelity latent-space mapping. **(A)** *Validation loss decreases with increasing training data size.* The validation MSE for the parameter-mapping model (Fig. 3) is plotted as a function of the total training dataset size (log scale). Each point represents the best-performing model for a given dataset size. **(B)** *Prediction quality improves with increasing training data size.* Representative test-set examples (rows) illustrate predictions obtained from models trained on progressively larger datasets (columns), compared with the corresponding inputs (left) and ground-truth outputs (right). High-quality structural predictions emerge at ∼3×10³ samples and saturate by ∼3×10 samples. **(C)** *Rotational augmentation enables strong performance with far fewer unique samples.* When each unique pattern is augmented by rotations to produce a constant total of 40,000 samples, the validation loss decreases sharply even for small numbers of unique samples. **(D)** *Non-blurred structured predictions with as few as ∼80 unique samples.* Test-set predictions from models trained on augmented datasets show that non-blurred, structurally accurate mappings emerge with ∼80 unique samples, demonstrating the effectiveness of rotational augmentation in reducing data requirements.

### Simulation-conditioned diffusion generates experimentally realistic bacterial patterns

Having established that SD VAE latents provide useful representations of both simulated and experimental patterns, and that rotational augmentation can substantially reduce the number of unique paired samples required for high-fidelity translation, we next applied this framework to simulation-to-experiment mapping. The goal of this stage was therefore not only to make simulations look more like experiments, but to learn an observation layer that preserves the mechanistic dependence of each pattern on its initial seeding configuration while restoring experimental morphology and replicate variability.

We collected ∼500 patterns and selected ∼400 for the training and validation, including replicates for each seeding configuration to capture biological variability. Swarming agar plates were prepared freshly each day^11^ under defined nutrient and agar conditions (Methods) and seeded with 0.1 μL of *P. aeruginosa* (PA14) cultures during exponential phase (OD600 = 0.2). Plates were imaged after 20 hours using a plate imager. Because patterns form within a circular plate, they are rotationally invariant. To this end, we augmented each simulation-experiment pair by multiple rotations, expanding the dataset to ∼40,000 image pairs. Before data augmentation, experimental images were processed for rotational alignment, contrast and brightness adjustment, and cropping (**Supplementary Figure 11**).

We first attempted to compress both simulated and experimental patterns into SD VAE latents and train a ResNet to map between them. However, the deterministic ResNet output could not capture the variability across experimental replicates, producing blurred images that resemble the averages of experimental replicates (**Supplementary Figure 12**). While confirming that SD VAE could represent both simulated and experimental patterns, in a manner that allows downstream mapping, the analysis indicated that a deterministic mapping was insufficient to capture the fine, local details present in experimental patterns.

To capture the variability in experimental patterns, we turned to diffusion models, which have emerged as effective tools for generating realistic images with inherent variability^51,69–72^. These models generate images from random Gaussian noise by an iterative denoising process, typically guided by a text prompt. In our case, the goal was to guide the denoising process with the simulated patterns. To this end, we used ControlNet, a trainable adapter attached to the Stable Diffusion’s denoising U-Net, which enables conditioning on image inputs, such as sketches, edges, and human poses^73^. Here, the simulated pattern served as the spatial “blueprint” for generating the corresponding experimental pattern. As in our earlier approaches, the ControlNet operated in the latent space, with the SD decoder reconstructing the final high-dimensional image.

We trained ControlNet on the augmented dataset (∼40,000 simulation-experimental pairs), using simulated patterns as conditioning inputs and experimental patterns as target outputs (**Figure 5a**, **Supplementary Figures 13 & 14**). During inference on a held-out test set (∼100 samples), the ControlNet generated experimental-like patterns from seeding configurations via the simulation-based blueprint (**Figure 5b**). Because the mapping is probabilistic, the model reproduced fine-scale experimental features, such as colony color gradients (reflecting cell density changes and pigment production), variations in branch number, and differences in branch orientation or length, while preserving the global structures imposed by the coarse-grained simulations.

**Figure 5:**
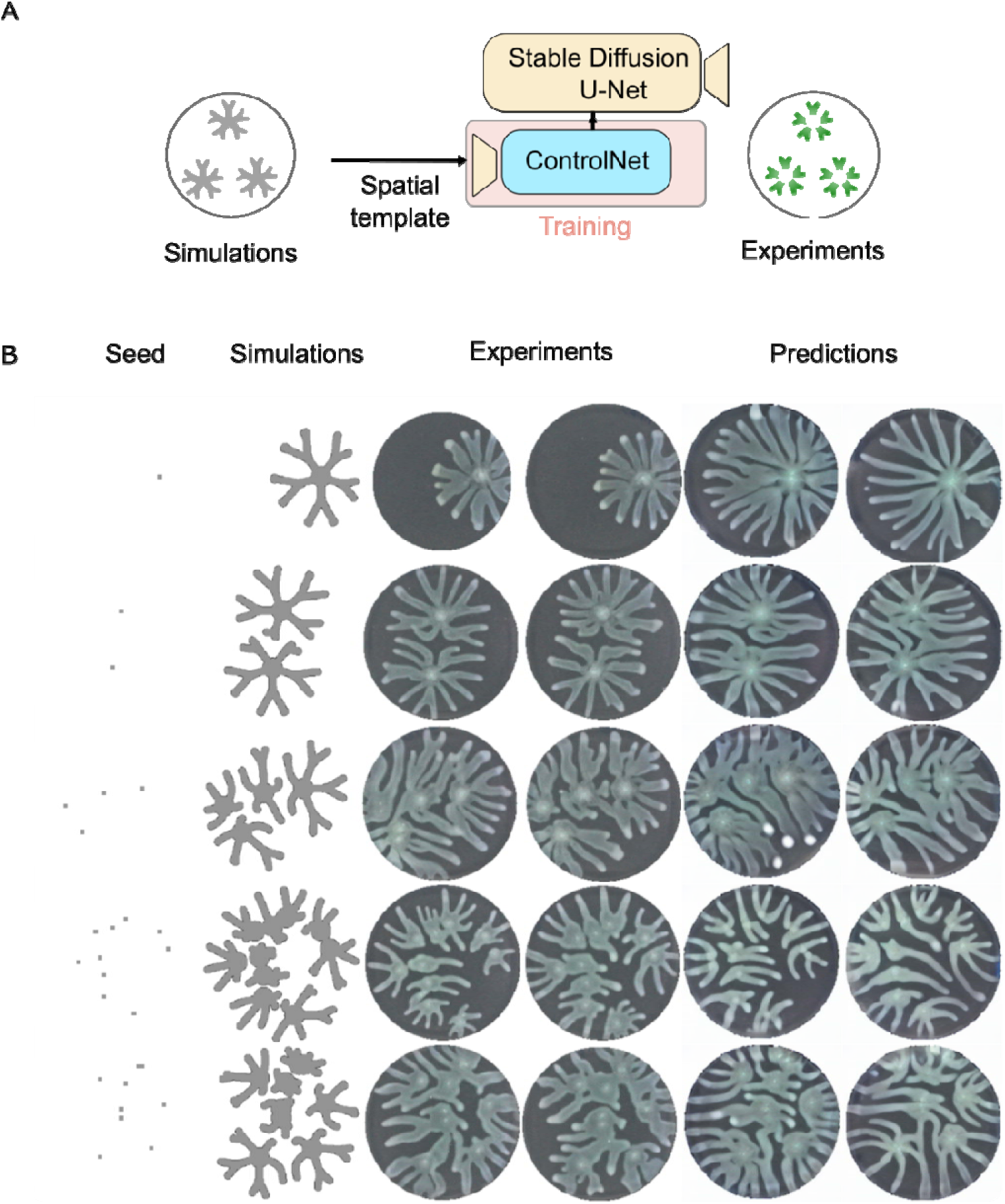
Simulation-conditioned diffusion generates experimentally realistic P. aeruginosa colony patterns. **(A)** *ControlNet framework for simulation-to-experiment mapping.* A ControlNet module attached to the Stable Diffusion U-Net is trained to generate experimental-like colony patterns conditioned on simulated patterns. Paired simulation–experiment images from diverse seeding configurations, including experimental replicates, are used for training. The simulated pattern provides a spatial template, while the diffusion model introduces realistic visual and stochastic features observed in experiments. **(B)** *Experimentally realistic predictions capture replicate-level variability.* For an out-of-sample test set, the trained model produces multiple plausible experimental-like patterns for each simulated input. Predictions preserve the global structure enforced by the simulation while reproducing key experimental characteristics, including variation in branch number, extent, and orientation across replicates. The model thus captures both the mechanistic constraints and the inherent stochasticity of swarming pattern formation.

Using the seeding configurations as the template instead of the simulated patterns produced plausible but less simulation-grounded predictions (**Supplementary Figure 15**), indicating that using simulations as the spatial template aids in pattern prediction. We also tested the influence of the spatial simulation template on the pattern prediction process by ControlNet, alongside other model parameters that are tunable during inference (**Supplementary Figure 16**). Lastly, generating multiple patterns from the same input captured realistic variability across replicates, with less variation than across distinct seeding configurations, matching experimental trends (**Supplementary Figure 17**).

To quantify ControlNet performance, we compared ControlNet predictions to experimental images using multiple similarity metrics: Learned Perceptual Image Patch Similarity (LPIPS)^74^, SSIM, Oriented Fast and Rotated Brief (ORB)^75^, and Maximum Mean Discrepancy distance using CLIP embeddings^76^ (CMMD) (see **Methods** for details). Across most perceptual and image-level measures, ControlNet predictions were closer to experiments than were PDE simulations(**Supplementary Table 1**). A stricter binary structural comparison gave a more heterogeneous result: for about one third of images, predictions were closer than simulations to the corresponding experiments (**Supplementary Figure 18**). This result is consistent with the goal of the model, which is not to exactly reproduce each experimental replicate, but to generate experimentally realistic patterns that preserve mechanistically relevant spatial structure.

Overall, these quantitative analyses corroborate our qualitative conclusion that ControlNet generated simulation-conditioned, experimentally realistic patterns. However, visual and image-similarity metrics alone do not establish whether the generated patterns preserve biologically useful information. We therefore next tested whether these generated patterns could serve as synthetic training data for inference on real experimental colony images.

### Simulation-conditioned synthetic data support synthetic-to-real inference on experimental patterns

Having established that simulation-conditioned diffusion can generate experimental-like bacterial patterns, we next asked whether these patterns preserve information that is useful for real biological inference. This provides a stronger test than image realism alone. In the forward direction, different initial seeding configurations generate distinct simulated and experimental branching patterns. The inverse problem asks whether the initial seeding configuration can be inferred from the morphology of an observed experimental pattern (**Figure 6a**). More broadly, microbial spatial patterns can act as records of initial conditions, environmental inputs, or growth histories^44,77,78^, making inverse inference from morphology a biologically meaningful problem rather than only a computational benchmark. Previous studies have shown that noisy simulated biological patterns can support distributed information encoding-decoding when large replicate datasets are available ^79^. Here, we asked whether a comparable inverse task could be performed on real experimental patterns, where experimental data are scarce. Our hybrid framework addresses this limitation by generating large numbers of simulation-conditioned, experimental-like patterns for defined seeding configurations.

**Figure 6:**
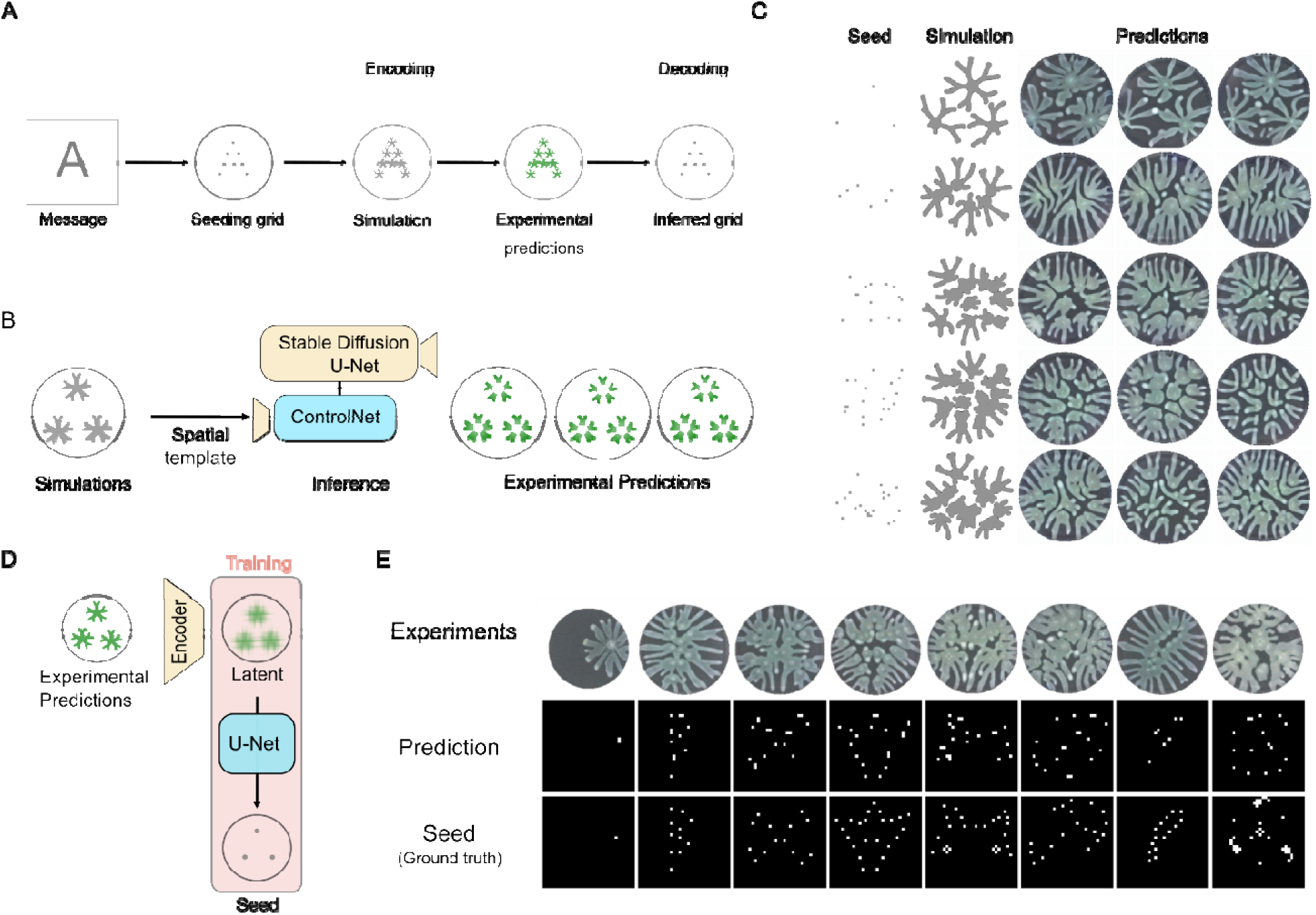
Simulation-conditioned synthetic data enable zero-shot inverse inference on real experimental patterns. **(A)** *Information encoding-decoding pipeline*. The spatial grid is used as a canvas for storing a message (letter A in this schematic). The message is broken down into discrete seeding positions, which are used as initial bacterial configurations for simulations. These simulations are then used as spatial templates to generate experimentally realistic, simulation-conditioned synthetic patterns using the trained model in Figure 5. All of the previous processes can be described as information encoding. A separate neural network is then trained on the inverse problem of decoding the initial positions from the predictions. **(B)** *Creation of a synthetic patterning dataset:* We used the simulation dataset used in Figure 2, as initial templates to the trained ControlNet framework shown in Figure 5, to generate 3 samples per simulation. A total of 30,000 input simulations result in a total output dataset size of 90,000 samples. **(C)** *Sample images of the synthetic patterning dataset:* Varying (rows) initial seeding conditions (first column) result in different simulated branching patterns (second column). Simulated patterns are then used as templates to generate multiple simulation-conditioned synthetic patterns using the trained ControlNet framework. **(D)** *Training a decoding model on synthetic data:* Synthetic patterns were compressed to their latent representations using SD VAE and a U-Net was trained to map from the latent patterns to the initial seeding configurations. **(E)** *Inferring initial seeding configurations from experimental patterns*: The performance of the trained model was judged on the test set consisting of real experimental patterns (first row). The predicted initial configurations (middle row) were close to the actual seeding configurations (ground truth, bottom row). The last two columns illustrate some of the limitations of the trained model on out of distribution samples, either when the seeds used for the patterns are very dense or the total seed count is more than those used in the training dataset.

To construct this test, we generated simulations from random seeding configurations and used each simulation as input to the trained ControlNet framework to generate multiple synthetic experimental-like samples per initial condition (**Figure 6b**). These generated samples were designed to capture experimental stochasticity in the branching process while preserving the mechanistic relationship between the initial seed configuration and the resulting colony morphology. The generated dataset included realistic variation in branch number, branch length, branch orientation, and color (**Figure 6c**). We then used this purely synthetic dataset, consisting of 90,000 samples generated from 30,000 seeding configurations with three replicates per configuration, to train a U-Net decoder (**Supplementary Figure 19**). The decoder mapped SD VAE latent representations of synthetic patterns to the corresponding initial seeding configurations (**Figure 6d**). Importantly, both training and validation used only generated synthetic patterns; the held-out test set consisted entirely of real experimental colony patterns that start from random or defined seeding conditions and was evaluated without fine-tuning.

When applied directly to real experimental patterns, the synthetic-trained decoder recovered the general locations of initial seeding positions across a range of test patterns (**Figure 6e**). This zero-shot transfer indicates that the generated patterns preserved information linking colony morphology to initial seeding configuration. Performance was less stable for denser seed arrangements or seed counts beyond those represented in the training distribution, indicating a boundary condition for out-of-distribution generalization. Quantitative evaluation using precision, recall, and F1 score with a four-pixel tolerance around each seeding point confirmed that the decoder trained exclusively on simulation-conditioned synthetic patterns recovered seed locations from real experimental images substantially better than the raw-simulation baseline. To test whether the gain comes from simulation-conditioned synthetic data rather than from the simulator alone, we trained a baseline decoder on raw simulations. On real experimental images, the decoder trained exclusively on simulation-conditioned synthetic patterns achieved a tolerance F1 score of 0.723 ± 0.208, compared with 0.218 ± 0.224 for the raw-simulation baseline. Precision and recall similarly improved from 0.377 ± 0.365 and 0.190 ± 0.235 for the raw-simulation baseline to 0.799 ± 0.206 and 0.720 ± 0.266 for the synthetic-trained decoder, respectively (Supplementary Table 2). Although the raw-simulation baseline model performed well on simulated pattern-to-seed mapping, its performance did not transfer to real experimental patterns without fine-tuning (**Supplementary Figure 20, Supplementary Table 2**).

Together, these results show that experimentally realistic generated patterns can reduce dependence on large experimental training datasets for inverse inference. More importantly, they establish task-based utility as a functional criterion for simulation-conditioned generation: the generated patterns are useful because they preserve morphology-linked information needed for inference on real biological images, not merely because they resemble experimental patterns visually.

## Discussion

This study establishes that mechanistic simulations can serve as spatial conditioning priors for generative modeling, and that the resulting synthetic patterns can preserve information sufficient for synthetic-to-real inverse inference on bacterial colonies. By combining a coarse-grained PDE model^15^ with foundation-model latent representations^51^ and conditional diffusion^73^, the framework converts mechanistic simulations that capture global colony structure into experimentally realistic pattern images that retain morphology-linked information for downstream inference. This complementarity — mechanistic structure from the simulator and fine-scale realism from the generative model — is the central design principle of the framework.

Self-organized pattern formation emerges from the interplay of biophysical processes operating at multiple scales. By construction, mechanistic models capture these processes and provide interpretability and predictive structure^2^, but they necessarily omit many of the microscopic features that give the visual richness observed in experiments^13,15,29,42^. Conversely, modern generative models excel at rendering high-fidelity images but lack mechanistic grounding and can fail to generalize out of the training data^80–82^. Our study shows that these two modeling approaches can be combined in a complementary way: coarse mechanistic simulations can serve as spatial conditioning priors for generative modeling, enabling experimentally realistic biological patterns that preserve global structure, restore fine-scale morphology, and capture replicate-level variability.

A central challenge in biological pattern prediction is that mechanistic models do not encode many of the heterogeneous and stochastic processes, such as local differences in motility^29^, surface wetness^30^, and pigment production^83^, that shape the experimentally observed morphology of real colonies. Our approach overcomes this limitation: the PDE simulation provides a spatial scaffold, whereas the conditional diffusion model learns the distribution of experimental-like morphologies around that scaffold. This division of labor allows each modeling component to operate in its domain of strength: the simulator enforces large-scale structure, while the ML model supplies the fine-scale realism that mechanistic models cannot capture. Importantly, this mapping could be learned with a modest experimental dataset: ∼400 experimental patterns, expanded by rotational augmentation to ∼40,000 paired examples, were sufficient to train the simulation-to-experiment generator. This result suggests a practical strategy for low-throughput biological systems, in which limited experimental data can be used to learn an observation layer that converts coarse simulations into experimentally realistic synthetic pattern data.

Our approach is distinct from commonly used scientific machine-learning strategies. Existing approaches often constrain neural network training with physical or biological constraints^47,82,84^, learn solution operators from simulated data^85,86^, infer parameters, governing equations or missing terms^46,87–89^ or accelerate dynamical prediction^17,25,27^. By contrast, our strategy does not seek to learn or replace the governing model itself. Instead, it uses the mechanistic simulation as a spatial conditioning input and learns an experimentally realistic observation layer on top of the coarse simulated pattern. This separation provides both conceptual clarity and modularity: the mechanistic simulator, foundation image model, and conditional mapping architecture can in principle be replaced independently without redesigning the entire pipeline. Thus, the framework complements existing scientific ML approaches by addressing a different problem: not how to infer or accelerate a mechanistic model, but how to convert its coarse output into biologically realistic synthetic observations.

This strategy also has boundary conditions. It requires a mechanistic simulator that captures enough global structure to serve as a meaningful spatial scaffold. If the simulator is qualitatively misspecified, a conditional generator may still produce realistic-looking images, but those images would have limited scientific value. The approach also depends on proximity between the experimental regime used for training and the regime used for deployment, as reflected by reduced inverse-inference performance for denser or otherwise out-of-distribution seeding configurations. Thus, simulation-conditioned generation should be viewed as a way to extend useful mechanistic models into experimentally realistic observation space, not as a substitute for mechanistic validity.

These results establish simulation-conditioned generative modeling as a framework in which mechanistic simulations provide structural validity, foundation image models provide expressive representations, and conditional generation links coarse simulations to experimentally realistic pattern data. The synthetic-to-real inverse task provides a functional validation of this framework. A decoder trained only on simulation-conditioned synthetic patterns inferred initial seeding configurations from real experimental patterns without fine-tuning, whereas a model trained on raw simulations did not transfer comparably. This result shows that the generated patterns preserve morphology-linked information relevant to biological inference, rather than merely matching superficial image statistics. In this sense, the framework converts a coarse mechanistic simulator into a source of scientifically structured synthetic training data for real experimental analysis.

This hybrid strategy offers a new path for predictive modeling in systems where mechanistic understanding and experimental fidelity are both indispensable. In biology, simulation-conditioned generative models could serve as hypothesis-testing tools for asking how global pattern structure and local morphological variability respond to genetic, environmental, or spatial perturbations. Because experimentally realistic morphology can be essential for downstream tasks, including strain identification^37,38^, morphotype classification^39,40^, perturbation inference^31,36^, or synthetic pattern design^21,42,44^, our model can act as a biologically grounded synthetic data generator. By coupling mechanistic simulations to experimental-like pattern generation, the approach makes it possible to create large, structured training datasets while preserving mechanistic relationships between inputs, spatial organization, and morphology-linked outputs.

The same logic may extend to other biological systems in which mechanistic models capture coarse spatial structure but fail to reproduce the experimentally observed morphology, including tissue morphogenesis^9^, biofilm development^90^, and microbial community assembly^78^. In each case, the relevant goal is not image realism alone, but the need to generate synthetic observations that are both realistic enough for downstream models and constrained enough to preserve the structure of the underlying system. More broadly, our work illustrates how foundation image models can extend mechanistic modeling beyond coarse simulation, offering a practical strategy for generating experimentally realistic synthetic observations of complex spatial phenomena across biology and the physical sciences.

## Methods

### Generation of randomized initial-seed/final-pattern training pairs

We used a coarse-grained optimization-based model of 2D branching pattern formation that was described in an earlier study^15^. The simplified model describes the branching dynamics by capturing cell growth, cell movement, nutrient uptake, and diffusion. The patterns are described using a branch width (W), branch density (D), where the local density in a branch is defined as 1/d, d denoting the distance of a branch tip with the nearest neighbor. The equations are given by:

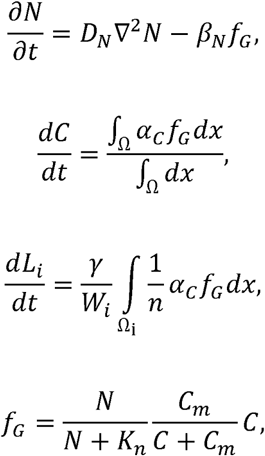

where the variable N denotes the nutrient concentration, C is the cell density per unit area of the colony, L denotes the branch length. Variables of this system of equations include *D_N_* which is the nutrient diffusivity, *β_N_* denotes the nutrient consumption rate, *f_G_* denotes the growth function, Ω denotes the growth domain, *γ* denotes the efficiency of colony expansion, W denotes the branch width, *α_C_* denotes the growth rate, Kn and Cm stand for half saturation nutrient constant and half saturation cell density constant respectively. Lastly, *i* stands for the i^th^ branch and *n* indicates the number of branches in the colony. The base parameter set for the simulations was:

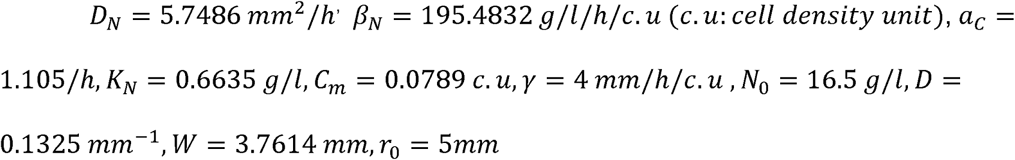

These parameter values were used to predict the standard experimental conditions of swarming patterns using 0.5% agar density and a high casamino acid concentration of 16g/L.

Figures that demonstrate mapping between two simulated parameter sets were constructed using the above parameter set, and a second set (which is referred to as thinner, denser branches in **Results)** with the following values:

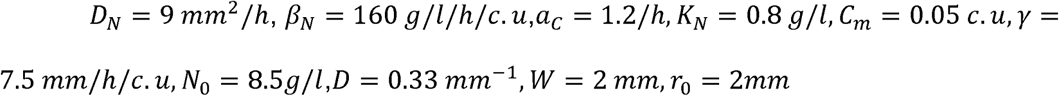

All the above simulations were run for a total time *t* =24 h.

The patterns were initialized with a circle at the center of radius *r*_0_ and branch bifurcation was triggered when the local density falls below a chosen threshold of 2/3*D*. Besides predicting the nature of the branching patterns at different environmental conditions, the model also predicted how the overall dynamics of branching would look like with the inclusion of multiple initial seeds. Here, instead of having a well-defined seeding grid, we chose randomized seeding configurations containing a variable number of seeds, with n uniformly sampled between 1 and 20. These seeds were placed randomly in a grid, within the zone where colony growth occurs.

The simulation was then run for a fixed amount of time, and the images of the initial seeds and final patterns were saved. We used the Duke Compute Cluster to run simulations in parallel such that there were in total around 100k initial seed, final pattern pairs.

### Using Stable Diffusion VAE to compress the final pattern to a lower dimensional representation

Instead of predicting the final patterns directly from the initial seeding configurations, we first mapped the final patterns to a low dimensional space. This mapping was done using a pre-trained VAE that has been trained for a text-based image generation model, Stable Diffusion^51^.

We used the Stable Diffusion VAE (SD VAE), which has been trained on a large-scale dataset, a curated version of LAION 5B known as LAION-2B-en^91^. Specifically, we used the Stable-Diffusion-v-1-5 library that was obtained from the diffusers library on Hugging Face. The VAE consisted of two components, an encoder that compressed high dimensional images to a latent representation and then reconstructed the images from this representation using the decoder.

The performance of SD VAE was first tested on the dataset consisting of simulated *P. aeruginosa* patterns in terms of reconstruction accuracy. SD VAE accepts an image that has 3 channels and compresses an image 48-fold. Simulated images had a dimension of 256×256×1 after minor image processing. The information was then triplicated across the 3 channels to generate a 256×256×3 image. SD VAE generated a latent representation of the image with dimensions 32×32×4. We used the SD VAE encoder to generate the latent representations of pattern images. These images served as inputs or outputs for various custom neural networks.

### Building a neural network emulator for pattern prediction

To predict the simulated patterns from different initial seed pairs, we trained a network to map from the initial seed to the latent representation of the patterns. We reasoned that mapping initial seeding configurations to low-dimensional latent representations would be easier than directly generating full-resolution images.

We used a custom dilated ResNet (**Supplementary** Figure 5) that was finetuned based on the performance of the network on the training/validation dataset. The architecture consisted of two blocks at the start and at the end of the architecture, first block converted the 3 channels in input into 64, and another (last block) converted 64 channels into 4 channels (to match the dimension of the latent representations obtained using the SD-VAE encoder.) In between these 2 layers, the network had 15 dilated ResBlocks. Each ResBlock had a total of 4 convolutional layers, with each layer having an increasing dilation rate—1,2,4 and 8. The outputs at the end of each convolutional layer were batch normalized and went through the ReLU activation function. Identity skip connections were implemented to accumulate the output before and after each block, allowing the network to avoid the vanishing and exploding gradients associated with deep neural networks. 30,000 samples (each being a pair of the initial seeding configuration and the latent representation of the final pattern) were used with a 90:10 train/validation split. We trained the network with early stopping implemented (with a patience of 70 and delta of 0.05) with 500 epochs, and an exponentially decaying learning rate starting from 5×10^-4^, Mean Square Error (MSE) loss and the Adam Optimizer algorithm for stochastic gradient descent.

The trained model was evaluated using an out-of-sample test set. This test set had a variable or fixed input grid, with the ground truth being the latent compression of the resulting simulated patterns corresponding to these seeds. The model performance on the test set was visualized by converting the output of the network, i.e. the predicted latent representation, back to the higher dimensional images using the SD VAE decoder. The absolute difference was taken (for individual pixels in the images) between the ground truth and the resultant decoded outputs of the network to calculate the absolute error between the two. SSIM scores were computed after normalization of images to the [0,1] intensity range.

### Testing feasibility of transfer learning

We implemented a pipeline that involved training a neural network to map between two different sets of patterns. First, we varied the parameters of the simulation to generate thinner, but denser branching, as specified in the first section of Methods. 30,000 initial seed-final image pairs were generated with this parameter set, and the latent representations of the final image pairs were obtained using the SD VAE encoder. The initial seed combinations chosen were the same as the ones corresponding to the default parameter set.

We then trained a new custom ResNet, which had the same architecture as described in the previous section albeit a difference in the first input layer of the network. Since the latent representations from the pre-trained VAE had 4 channels, we modified the first input block to convert 4 channels to 64 channels (instead of 3 to 64 conversion). The network was then trained to map between the latent representations of the simulated patterns from one parameter set to another. The trained model was tested on an out-of-sample test set. The model performance on the test set was visualized by converting the output of the network, which in this case was the latent representation of the simulated patterns corresponding to the second set, to the higher dimensional representation by using the SD VAE decoder.

We then tested the performance of mapping between the two different sets of patterns with exponential changes in the number of training images ranging from 100 to 12,800. To account for the possibility of data augmentation due to rotational invariance in the images, we also selected unique image samples ranging from 10 to 1600. Each sample size was then augmented to a total of 40,000, with datasets having less unique samples having a higher number of augmentation images per unique image. To test the effect of the training data size with and without augmentation, neural networks were trained in parallel that had the same architecture and fixed hyperparameters, with the only varying component being the datasets.

### Using diffusion models from Stable Diffusion and spatial conditioning using ControlNet

We modified the ControlNet implementation for spatial conditioning of diffusion models with simulated patterning conditions, together with the Stable Diffusion checkpoint v1-5 from Huggingface. The model was trained on 40,900 simulated- experimental patterning pairs, with simulated images being used as an spatial conditioning inputs to the model and experimental images as desired model outputs, and blank text was used as the input text conditioning.

The model was trained for 5 epochs with a batch size of 4 and learning rate of 10^-5^, which took approximately 12 hours on a single A5000 GPU (Duke Computing Cluster). Model inference was tested on 96 simulated-experimental patterning pairs. Inference parameters used were as follows, Prompt: “”, Image samples:2, Control strength:1, Guess mode: FALSE, DDIM Steps:50, Guidance scale: 15.1, Seed: 729397049, Eta:0, Added prompt: “”, Negative prompt: “longbody, lowres, bad anatomy, cropped, worst quality, low quality”. These generic negative prompts were inherited from the default ControlNet inference workflow and were kept fixed across all reported inference runs. To test the baseline effect of the diffusion models, we trained another ControlNet with input seed- experimental patterning pairs with all other hyperparameters of the network kept fixed. We observed that the prediction performance could also be controlled to a great degree during model inference, and the effects of different control knobs were tested in the ablation study. These were:

1. Guess mode parameter, which controlled the level of effect of the initial input spatial condition to the final patterns
2. Removing negative prompts, which are used in the classifier-free guidance (CFG) scaling, that guides the denoising process in Stable Diffusion during inference.
3. Addition of default positive prompts listed in the original ControlNet implementation
4. Effect of lower(0.85) and higher (1.25) conditioning control strength
5. Higher (50) DDIM sampling steps
6. Lower guidance scale (9.0), listed in the original ControlNet implementation and also used for model training.

The stability of the prediction from the model was also verified: the model generated plausible predictions across different random seeds controlling the starting random noise for the diffusion process.

### Experimental data generation using MANTIS liquid handler

All the experiments in this study were done with *P. aeruginosa* PA14. Plates were streaked with glycerol stocks of PA14 and single colonies from the plates were used for overnight incubation in a shaking incubator at 37°C, 225 rpm for 16-20 hours. OD600 was measured the next day using a TECAN plate reader and transferred to a fresh LB tube at an OD600=0.05 and incubated at 37 °C for around 3 hours to allow for exponential growth. Swarming agar constituents were as follows: 1x Phosphate Buffer (5X buffer stock solution was prepared by dissolving 12 g Na_2_HPO_4_ (*anhydrous*), 15 g KH_2_PO_4_ (*anhydrous*), and 2.5 g NaCl in 1 L of water and was sterilized by vacuum filtration), 0.1mM CaCl_2_ (0.55 g CaCl_2_ in 50ml water was prepared as the 1000x stock solution, sterilized by vacuum filtration), 1mM MgSO_4_ (dissolving 6.02 g MgSO_4_ in 50 ml water was used as 1000x stock solution, sterilized by vacuum filtration), 0.5% agar (BD Difco™ 214530, agar was prepared freshly on the day of the experiment and sterilized by microwaving) and 16 g/L casamino acid(Gibco™ Bacto™ 223120, 200 g/L was used as the stock solution and kept at 4 °C for storage, sterilized by vacuum filtration). 20 ml swarming media was poured per Petri dish (100mm x 15 mm, Falcon), and the plates were allowed to cool down and dry with lids closed for an hour. PA14 cultures were adjusted to an OD ∼ 0.2 and agar plates were inoculated with 0.1 μL per seed using MANTIS liquid handler. After inoculation, the plates were dried with lids open for 15 minutes near a bench top flame and then were incubated upside down in a 37°C incubator for 20 hours.

Afterwards, the plates were imaged using UVP Colony Doc-It Imaging Station with epi white light with lids off in the right-side up position.

Random grids were designed for experiments that had a uniform distribution between 1 and 20 seeds per plate, with the seeds also being uniformly distributed throughout the circular plate area. 1536 well plate was chosen as the basis for the plate design, with the possible area further being restricted according to dimensions of the petri dishes. The highest possible resolution between two seeds corresponded to 1 well size in a standard 1536 well plate, which corresponded to a distance ∼ 2.25mm. Some of the designs for the plates also included a predetermined seeding configuration. Two to six biological and technical replicates were chosen for each seeding configuration to capture the experimental noise. Multiple batches of experiments were conducted, and after preprocessing which included manual rotation of some of the imaged patterns to align with the seeds or other technical replicates and filtering some of the outlier plates, a total of 409 final experimental patterns, including technical replicates, were obtained.

Experimental data was augmented by applying rotation of both the seed and the final experimental patterns around the center of the plate. The center of the plate was calculated by first identifying the circular boundary of the plate using Canny edge detection and then the center was determined using Hough circular transform. Each of the 409 experimental patterns were rotated by every 3.6 degrees to result in a 100x increase in the training/validation data to 40,900. These experimental images then went through some basic image processing tasks like cropping to remove plate boundaries.

Based on the experimental seeding grid, patterns corresponding to the simulations were also generated, with identical duplicate simulations serving as simulation counterparts for technical or biological replicates in experiments. The simulations were also rotated at similar angles as the experimental patterns to result in a total of 40,900 simulated patterns-experimental pattern pairs. Latent representation of these simulated and experimental patterns was generated, which served as inputs and outputs to the neural network respectively. Model performance was tested on ResNet and the ControlNet pipeline. The ControlNet pipeline does not need explicit latents as inputs and outputs to the model. Instead, the latents were computed internally during the training process, with a tiny network used for the encoding of the spatial template and the Stable Diffusion Encoder used for the experimental patterns, and the final outputs are decoded using the Stable Diffusion Decoder.

### Evaluation metrics

For model evaluation metrics, a test dataset was constructed with corresponding simulations, experiment and prediction triplets. A total of 96 images including replicates were chosen for each imaging modality. Since simulations are deterministic, the simulations were repeated for experimental replicates. Each seeding configuration had 2 experimental replicates, with a total of 48 unique conditions. For prediction images, the first two random seeds shown in Figure 5 were used to generate predictions corresponding to the simulation-experiment pairs in the test set. Evaluation metric testing was done on image triplets for LPIPS, SSIM and ORB scores, and done on entire datasets for CMMD. LPIPS (Learned Perceptual Image Patch Similarity) score reflects the perceptual similarity of images, calculated by comparing the latent distributions of the images using the VGG encoder. SSIM is used for structural similarity. ORB(Oriented FAST and rotated BRIEF) algorithm is used for local feature detection in computer vision tasks. CMMD, is an unbiased image estimator metric based on CLIP embedding of images and comparison of maximum mean discrepancy distance with the RBF kernel. It is similar to FID(Fréchet Inception Distance) for assessing the quality of generated images using a deep learning model, but is more aligned with human raters, and requires fewer images for a stable performance evaluation.

### Contrastive learning

A Siamese network was employed to quantify the dissimilarity between image pairs. The model consisted of convolutional neural networks that take pairs of images as input and are trained using a contrastive loss function. This loss makes the model produce similar embeddings for images generated under the same seeding conditions and dissimilar embeddings for images from different seedings. The training dataset consisted of simulated image pairs with binary labels: pairs from the same seeding condition were labeled as similar (i.e. 1), while those from different seedings were labeled as dissimilar (i.e. 0). A total of 96 image pairs were randomly assembled, in which half are “identical” and half are “different”. A random 80% of the dataset was used for training and 20% reserved for testing. Early stopping was applied to halt training if validation accuracy did not improve over 10 consecutive epochs.

The trained model outputs a dissimilarity score; higher scores indicate greater differences between patterns. We applied the model to unseen image pairs, including simulation-experiment and prediction-experiment comparisons, to evaluate the similarity between predicted or simulated patterns and their corresponding experimental results under the same conditions. To eliminate color as a confounding factor, all images were converted to black and white before training or testing.

### Computing the inverse problem for information encoding-decoding demonstration

U-Net architecture was employed for solving the inverse problem of mapping the initial seeding conditions as the output of the network from the experimental patterns as input. For the training of the network, we first used the simulation dataset used in Figure 2 that is different from the dataset used to train the ControlNet framework in Figure 5. This was done to demonstrate that ControlNet can generalize well on different datasets. Each simulation corresponded to a unique seeding configuration, and from each unique simulation, we generated 3 synthetic experimental-like patterns to act as biological replicates. For generating these synthetic patterns, we used the trained ControlNet with each sample generated from a random starting noisy seed, with rest of the inference parameters kept the same as one used in Figure 5. The total data size generated was 90,000 (30,000 unique simulations x 3 replicates) synthetic patterns, seed pairs. We then used this synthetic experimental-like dataset as the input to the network, with the respective seeding configurations from the paired simulations serving as the ground truth outputs. Also, the training pipeline first compressed the high dimensional images to latent representations (here the synthetic experimental-like patterns) using SDVAE Encoder (v 1-5, same as earlier figures) for efficient downstream training, before mapping them to the respective ground truth seeds. We used a combination of DICE loss and Binary Cross Entropy loss for training the network, with an extra weight given to positive samples to accommodate for the sparse seeding grid (weightage to DICE – 0.5, positive weight 2.0). Total trainable parameters in the U-Net were 505k, and the training/validation split was 90/10. The network was trained for a maximum of 500 epochs with early stopping and a learning rate of 5×10^-4^.

The performance of the network was tested on an out of sample test set that consisted solely of actual experimental samples, done without any fine-tuning of the network. For calculating the mean and standard deviation values used for estimating the Precision, Recall and F1 scores with tolerance (of 4-pixel radius, points matched using Hungarian Matching algorithm), the total dataset size was 132 images. These included the 96 images used as test set in Figure 5 and an additional 36 images with fixed seeding conditions having a mixture of in and out of distribution samples (additional number of seeds, more densely spaced seeds) compared to the training dataset.

As a baseline estimate, we tested the performance of mapping from simulation to seeds and then transferred to the experimental dataset. The simulation dataset used in Figure 2 was augmented with 3 rotations per image to result in a total dataset size of 90,000. The network was then first tested on the test set consisting of 107 simulation, seed pairs. Consequently, the performance was tested on the experimental test set consisting of 132 images.

## Acknowledgments

We thank Roarke Horstmeyer, Dongheon Lee, Yasa Baig and Jing-Mei Qian for helpful feedback, Helena Ma for initial setup and troubleshooting of MANTIS liquid handling system, Yasa Baig and Irida Shyti for discussions on model building, Tom Milledge and Duke Compute Cluster for assistance and resources on high performance computing, Eric Monsoon from the Center of Data and Visualization Sciences at Duke University Libraries and Zhengqing Zhou for assistance with figures. This work was partially supported by grants from the Office of Naval Research (LY: N00014-20-1-2121). The funders had no role in study design, data collection and analysis, decision to publish, or preparation of the manuscript.

## Author contributions

K.S. and L.Y. conceived the initial study. K.S built the simulation dataset along with guidance from L.Y. L.Y adopted the Stable Diffusion VAE on the simulation dataset and designed the initial ResNet models. K.S contributed to further computational model building and testing used in Figures 2,3 and 4 and adopting the ControlNet used in Figure 5 under guidance from L.Y. H.M.D and K.S contributed to the test of transfer learning used in Figure 4 under guidance from L.Y. K.S designed the experimental study used in Figure 5 with help from H.M.D and guidance from L.Y. K.S designed the information encoding-decoding pipeline in Figure 6 with guidance from L.Y. K.S, H.M.D, C.A.V. and A.H performed experiments. K.S and J.L conducted prediction model evaluation. J.L and E.S assisted with data interpretation. A.H and E.S assisted with checking the entire computational pipeline. K.S and L.Y wrote the manuscript with input from all the authors.

## Data availability

All data in the main text and the supplementary information are available at Huggingface Datasets: https://huggingface.co/datasets/HotshotGoku/Simulation_templated_pattern_prediction /

## Code availability

Code is currently being actively maintained at the youlab GitHub repo: https://github.com/youlab/Simulation_templated_pattern_prediction

