## Supplementary Materials for "Simulation-conditioned generative modeling for biologically realistic pattern prediction"

**List of supplemental figures and tables**

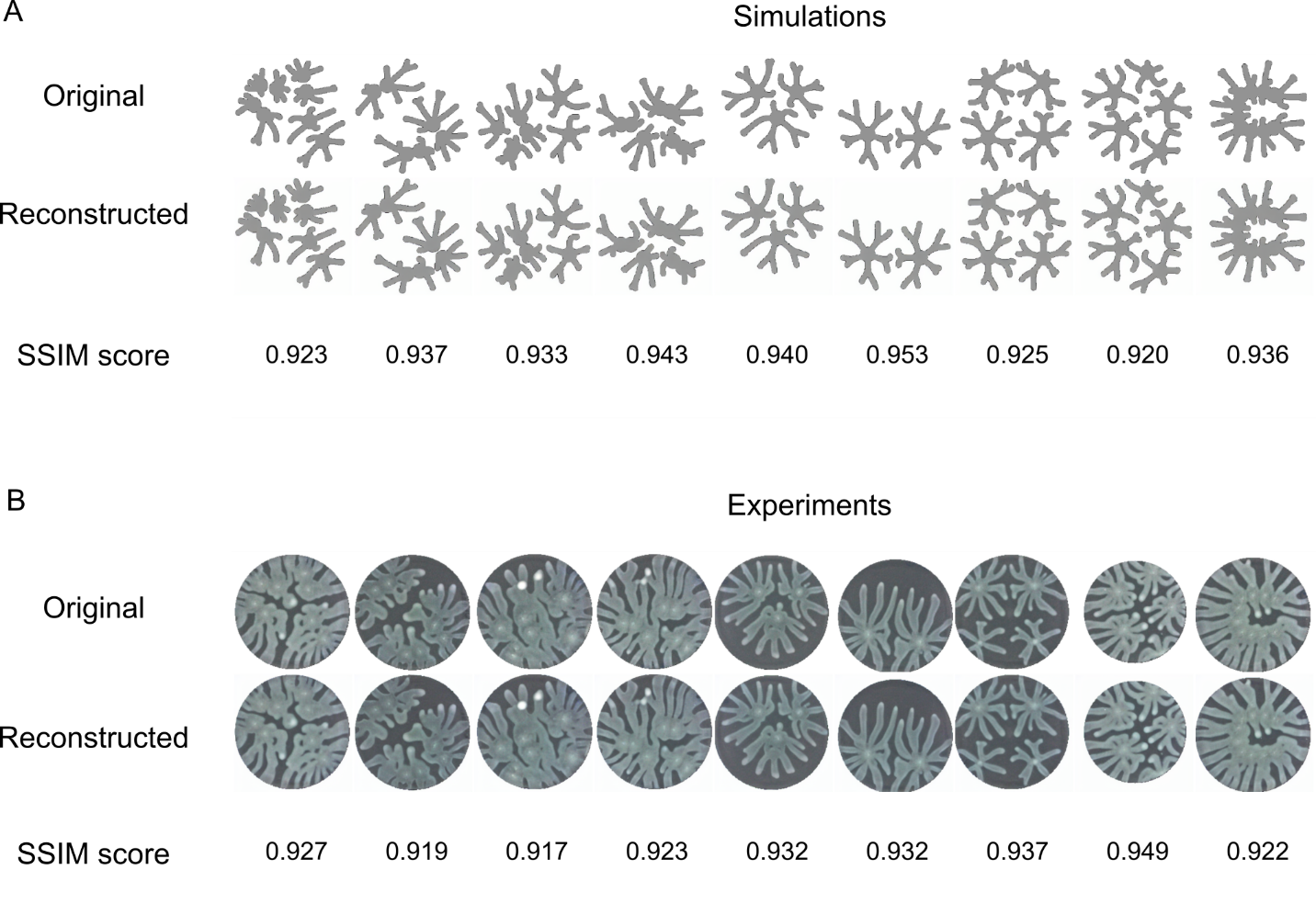

**Supplementary Figure 1: SSIM metrics confirm high-fidelity reconstruction of simulated and experimental patterns.**

(A) *Simulated patterns.* Representative simulated branching patterns (top) are shown alongside their reconstructions (middle) and corresponding SSIM scores (bottom). The examples span the diversity of the simulation dataset, including sparse random configurations and dense predefined designs such as the “C”-shaped pattern.
(B) *Experimental patterns.* Representative experimental *P. aeruginosa* branching patterns (top), their reconstructions (middle), and associated SSIM scores (bottom) are shown for the same seeding configurations as in panel A. Reconstruction fidelity is high for both datasets, with simulated patterns showing slightly higher SSIM scores on average.

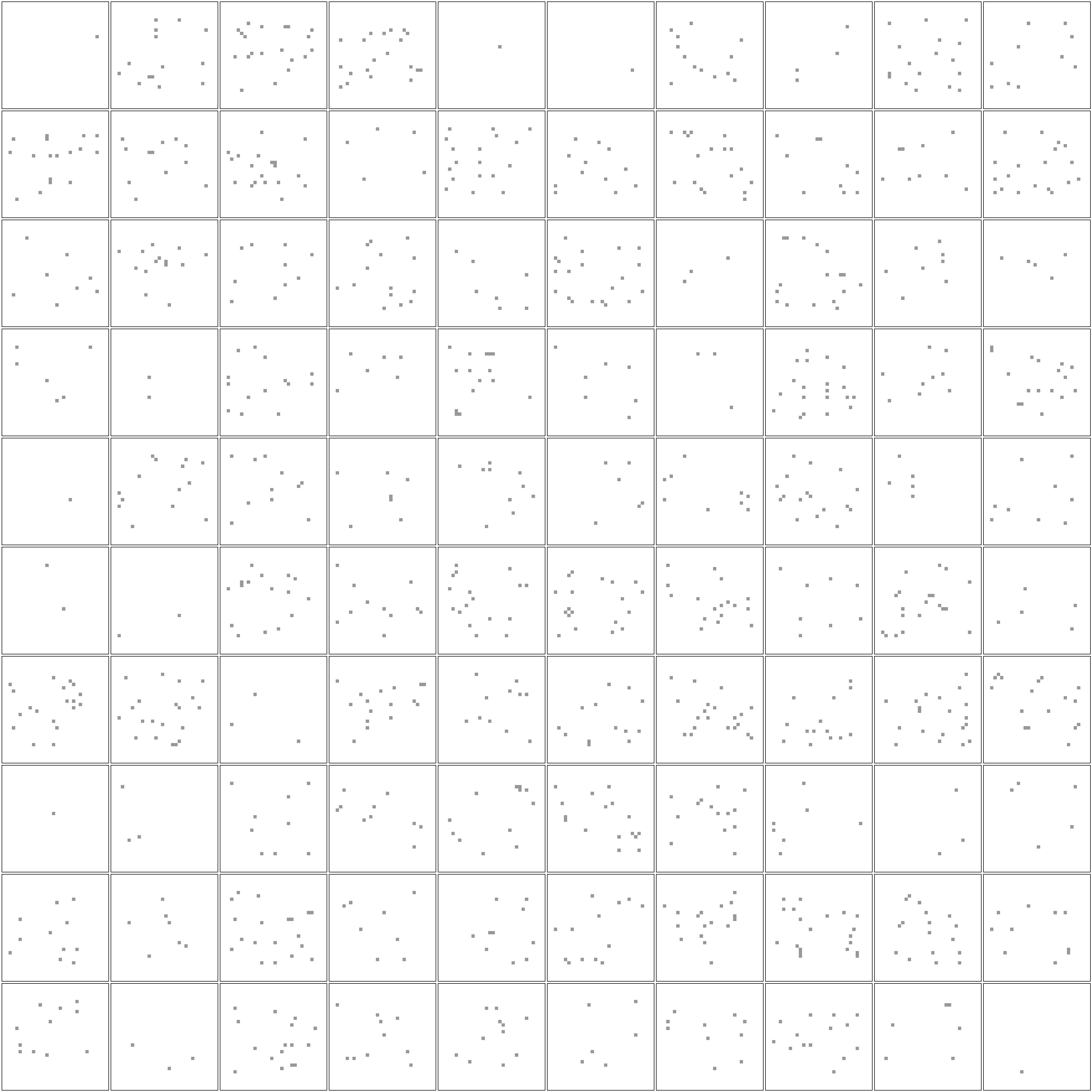

**Supplementary Figure 2: Sample seeding configurations for PDE simulations and inputs to the ResNet.**

Shown are 100 sample seeding configurations, each containing 1–20 seeds randomly distributed across the grid. These images serve as inputs to the ResNet model and have dimensions of 32 × 32.

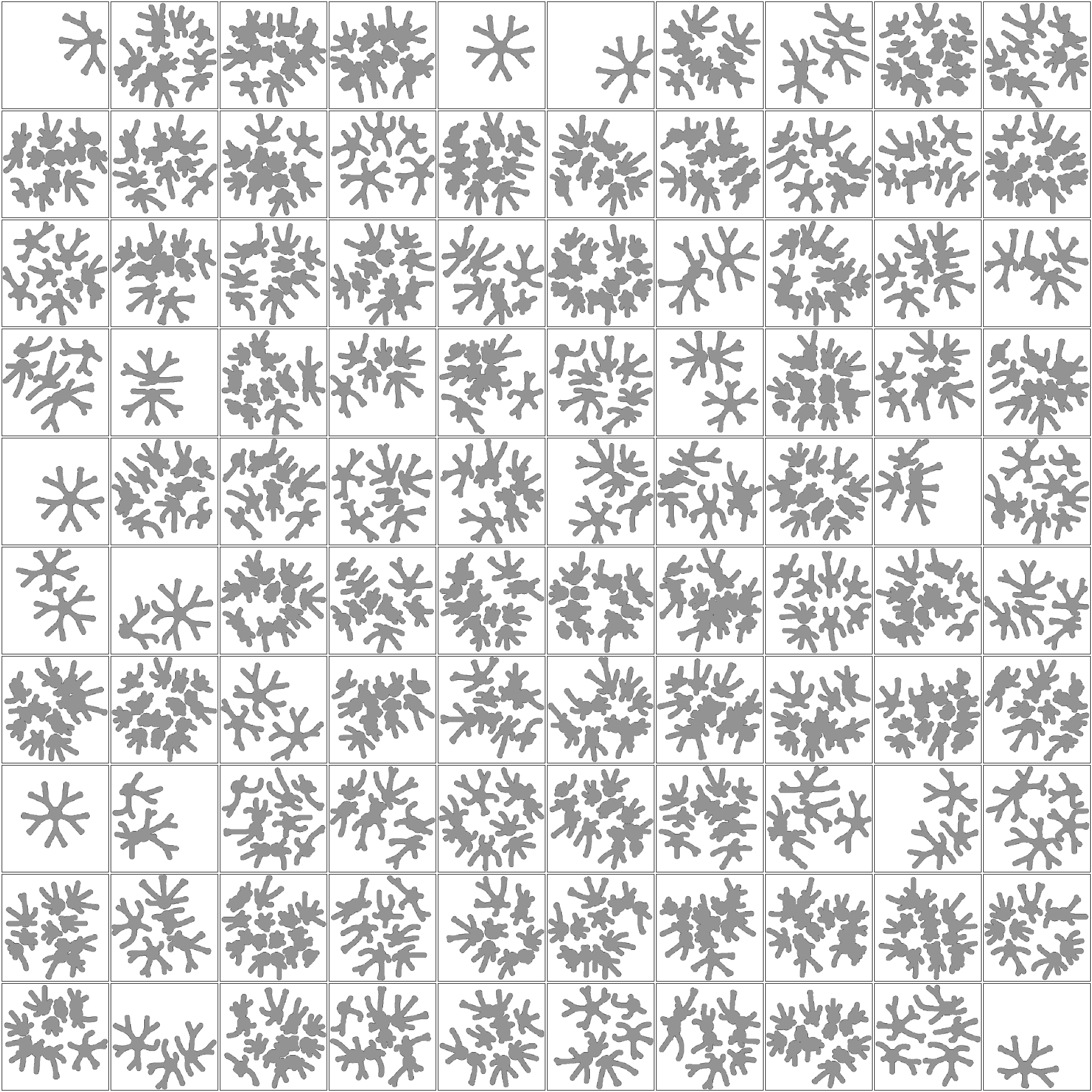

**Supplementary Figure 3: Simulated branching patterns from the seeding configurations shown in Supplementary Figure 2.**

A PDE model with a fixed parameter configuration (outlined in **Methods**) was used to simulate branching patterns from various seeding configurations. These images have dimensions of 256x256.

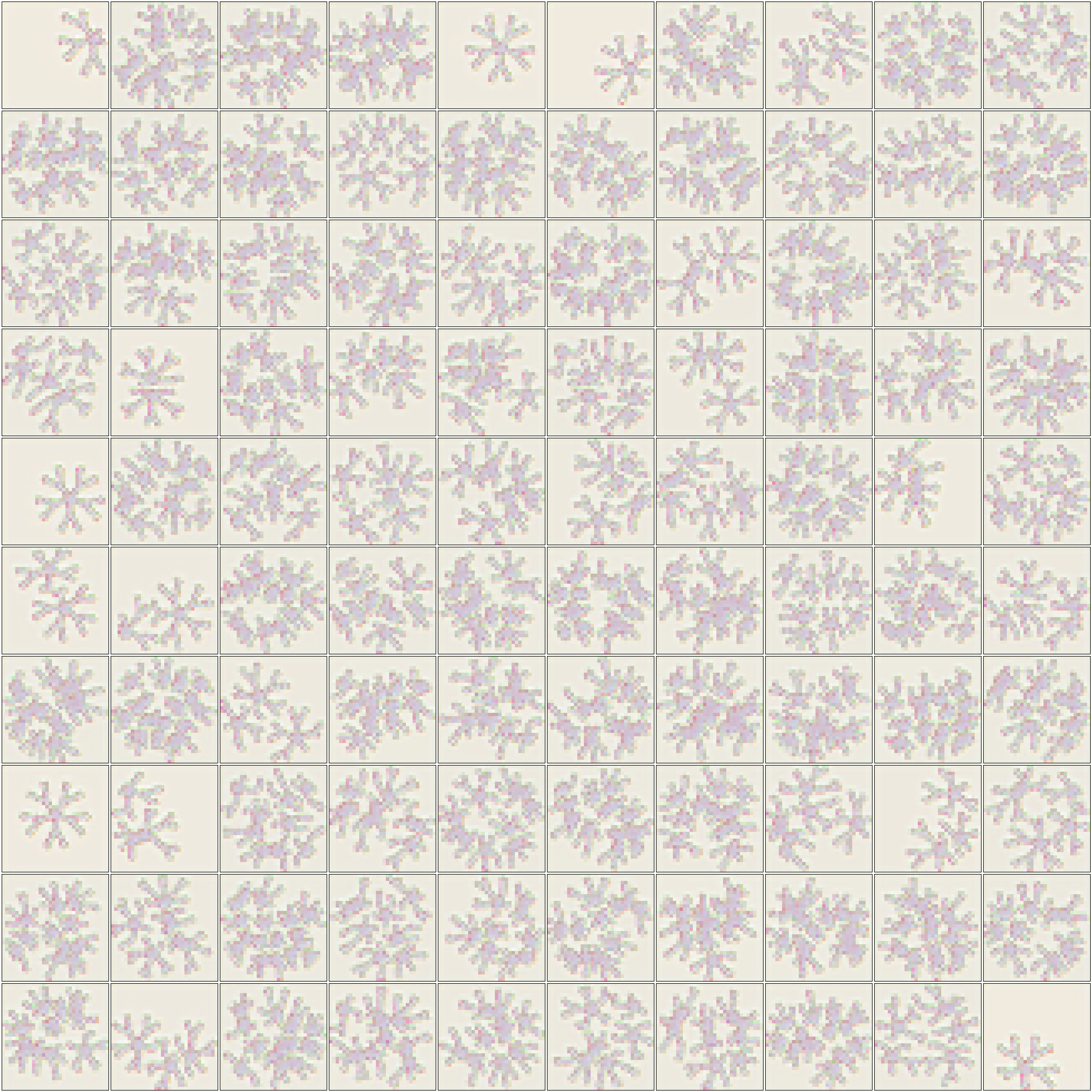

**Supplementary Figure 4: Visualization of the latents corresponding to the patterns in Supplementary Figure 3.**

Patterns in Supplementary Figure 3 are encoded into latents using the SD VAE encoder. These latent images have dimensions of 32 x 32 x 4. All 4 channels are used for visualization using an RGBA scheme.

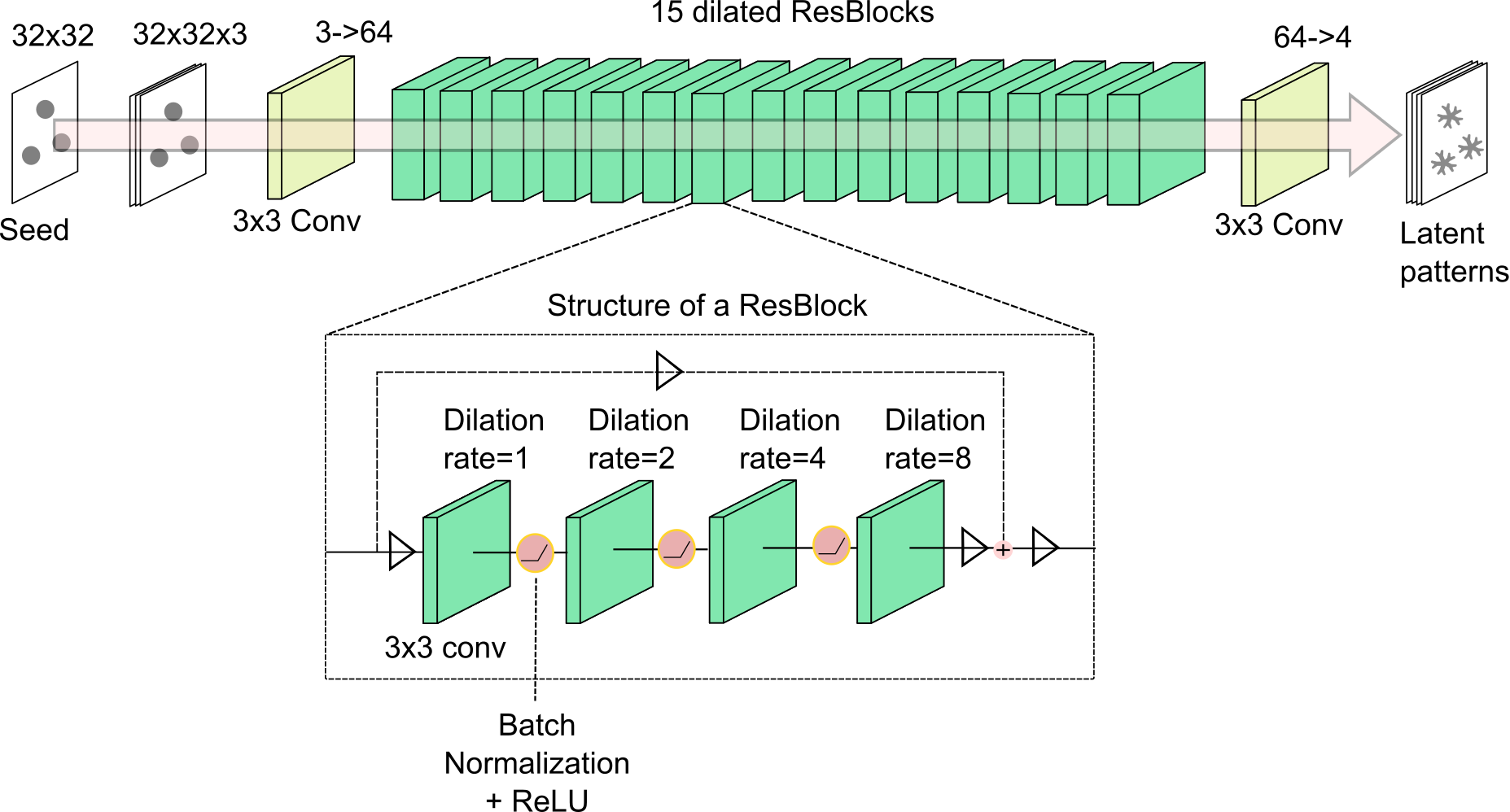

**Supplementary Figure 5: The ResNet architecture**.

The dilated ResNet accepts a 32 x 32 x 3 input image (seeding configurations, channels were triplicated) and outputs a 32 x 32 x 4 output image (corresponding to the latent representations). The architecture consists of 15 dilated ResBlocks (outlined in teal), with each block consisting of a series of convolutional layers with dilation rates of 1,2,4, and 8 (inset). A 3 × 3 convolutional layer at the beginning and another at the end convert the input and output to the desired channel dimensions. This architecture is used in Figure 2, with 3 additional ResBlocks for a total of 18 being used in Figures 3, 4 and Supplementary Figure 12.

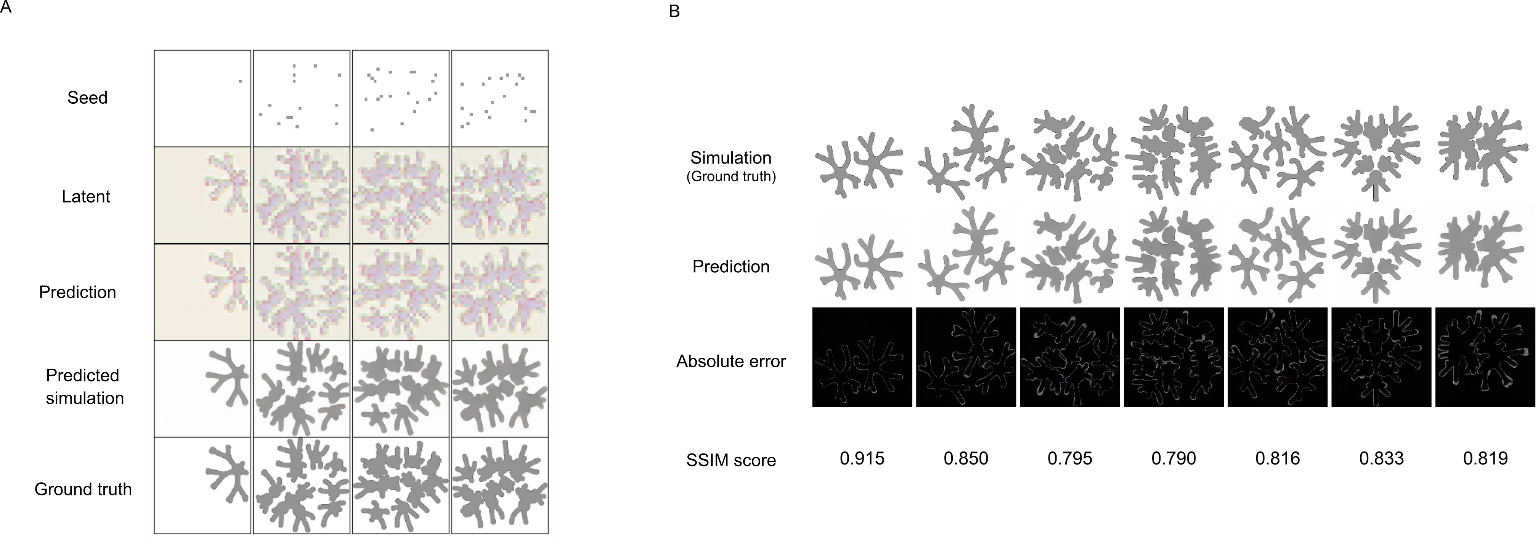

**Supplementary Figure 6**: **Performance of the ResNet on emulating the simulated patterns from initial seeding configurations**

(A) *Training samples:* Rows from top to bottom show the initial seeding configuration, ground-truth simulated latent representation, ResNet-predicted latent representation, decoded predicted pattern, and ground-truth simulated pattern.
(B) Model performance: *Absolute error of the prediction made on test set and ground truth simulated patterns*. Our model captures various local and global dynamics with high fidelity, with small scope of improvement at the colony boundaries.

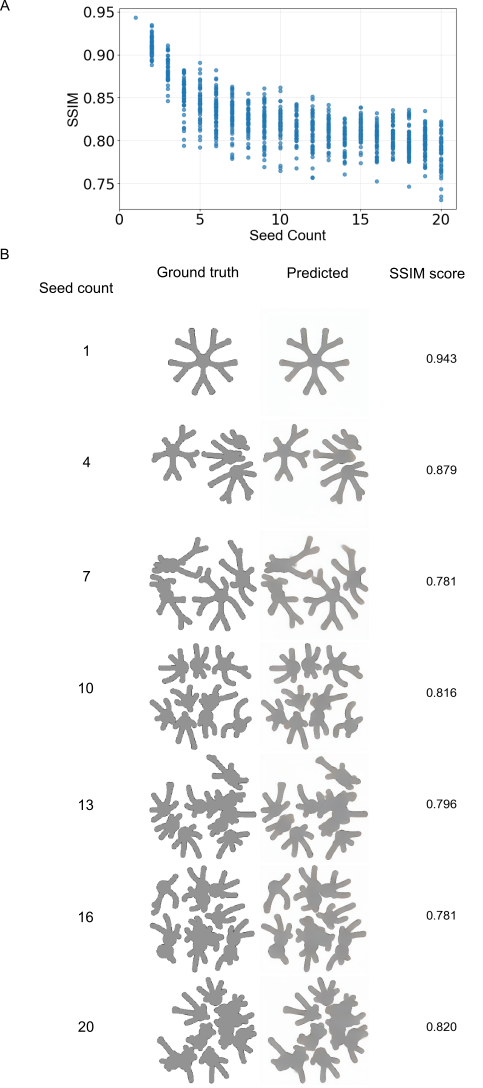

**Supplementary Figure 7**: **SSIM scores of our emulated model remain mostly constant throughout the range of seeding configurations**

(A) SSIM scores are compared for the emulated and the simulated images for 1000 random seeding configurations from the test dataset. The scores were relatively high, ~1 for images with a low number of seeds (<3) but levelled off at ~0.80 for the rest of the seeding numbers, including 20 seeds, the maximum used in this study.
(B) A sample of simulated and emulated patterns shown with an increasing number of seeds. Images having a lower number of seeds (1 seed, first row image) showed an SSIM score of 0.943, with the rest of the images having values close to 0.80 (mean = 0.824)

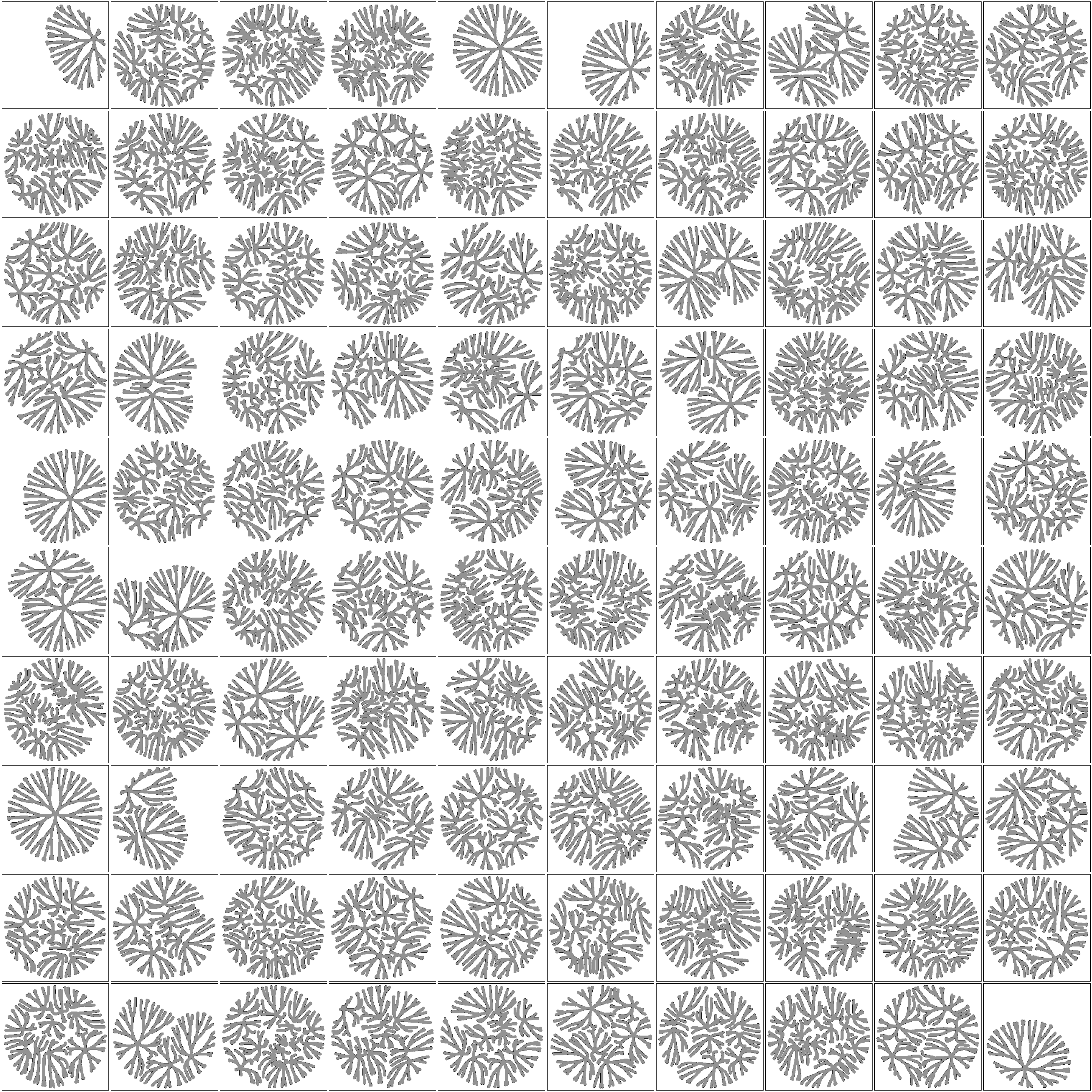

**Supplementary Figure 8**: **Sample of patterns simulated with same seeding configuration but different model parameters.**

The seeding configurations shown in Supplementary Figure 2 are used to simulate branching pattern formation using a different parameter configuration, which generates patterns with thinner and denser branching (outlined in **Methods**), and the end-point patterns are shown in the figure. Latent compression of these patterns served as outputs for our ResNet in Figure 3. These images have dimensions of 256x256.

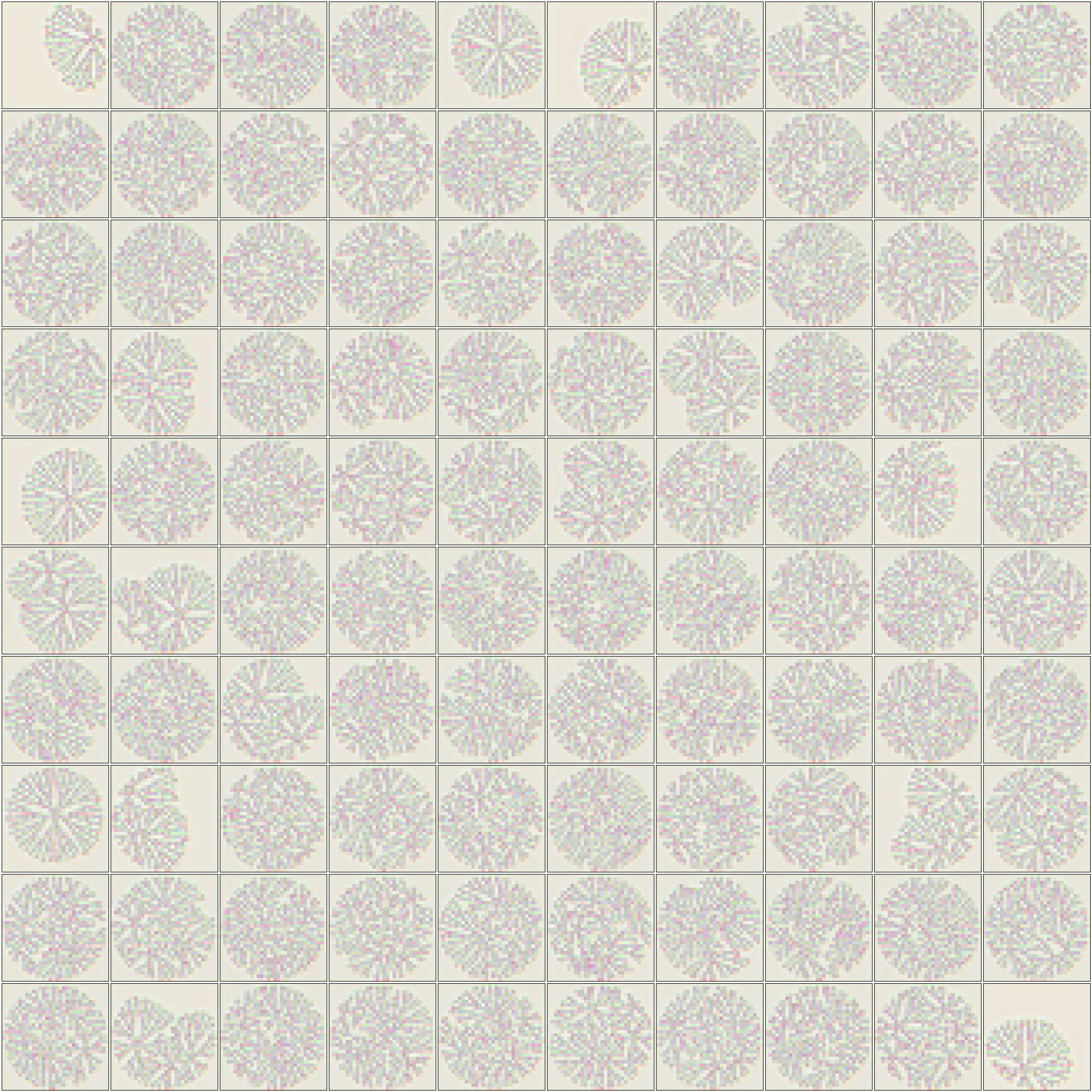

**Supplementary Figure 9**: **Latent representations of the images in Supplementary figure 8***.* Patterns in Supplementary Figure 8 are used to generate the latents using the SD VAE encoder. These latent images have dimensions of 32x32x4. All 4 channels are used for visualization using an RGBA scheme.

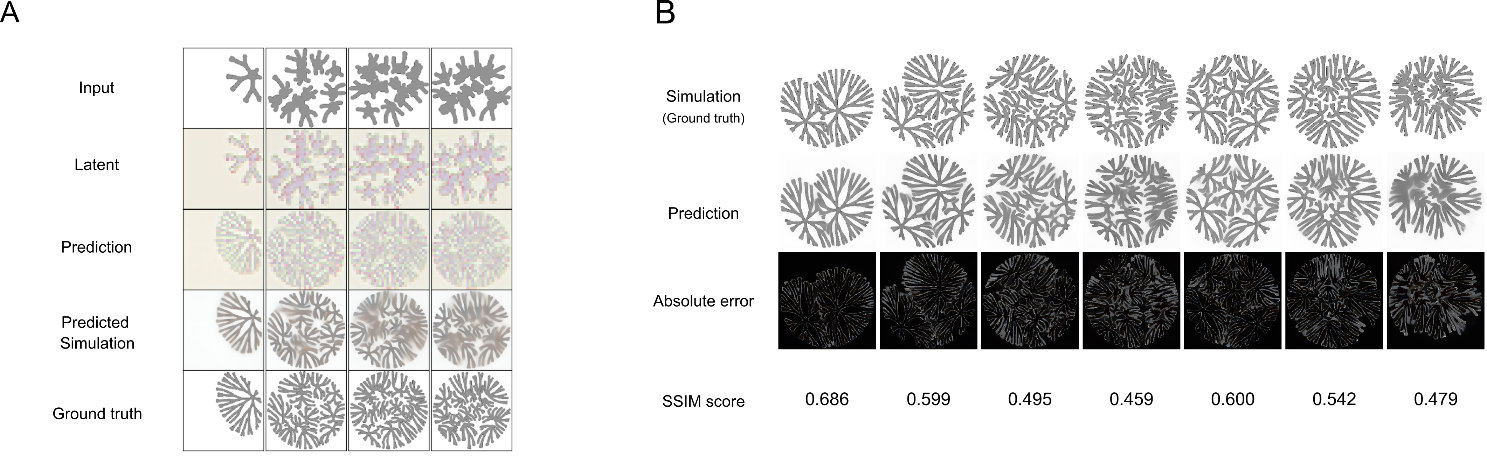

**Supplementary Figure 10**: **Performance of latent-space mapping using ResNet.**

(A) *Training samples*: Rows from top to bottom show patterns simulated with the first parameter set, ground-truth latents for the first parameter set, ResNet-predicted latents for the second parameter set, decoded predicted patterns for the second parameter set, and ground-truth patterns for the second parameter set.
(B) Model performance: *Absolute error of the prediction made on test set and ground truth simulated patterns*. Our model can capture various local and global dynamics, with some scope of improvement at the colony boundaries, and with more dense seeding patterns, as shown in the middle column seeding configuration.

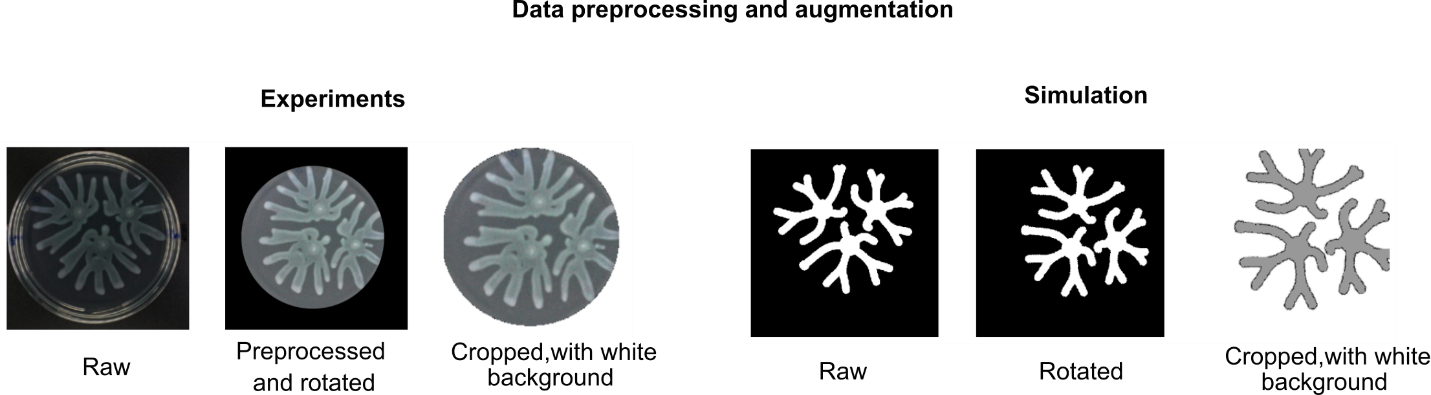

**Supplementary Figure 11**: **Experimental and simulation data preprocessing for analysis shown in Figure 5 and Figure 6**

Raw experimental images (outlined in **Methods**) were enhanced by improving the brightness and contrast of the image and applying a central mask. Simulations also went through a preprocessing step, an inverse mask is applied to the images such that black pixels correspond to the patterns, which are then converted into gray values (152 on the 0-255 scale) for better visualization. For data augmentation, each available experimental, simulation image pair was rotated by consecutive multiples of 3.6 degrees. The figure denotes a sample with a rotation of 45 degrees applied. Experimental rotations are made by identification of the circular boundary and then applying the rotation to each point in the circle. All the points inside an inscribed circle in the simulations are rotated by the chosen angle.

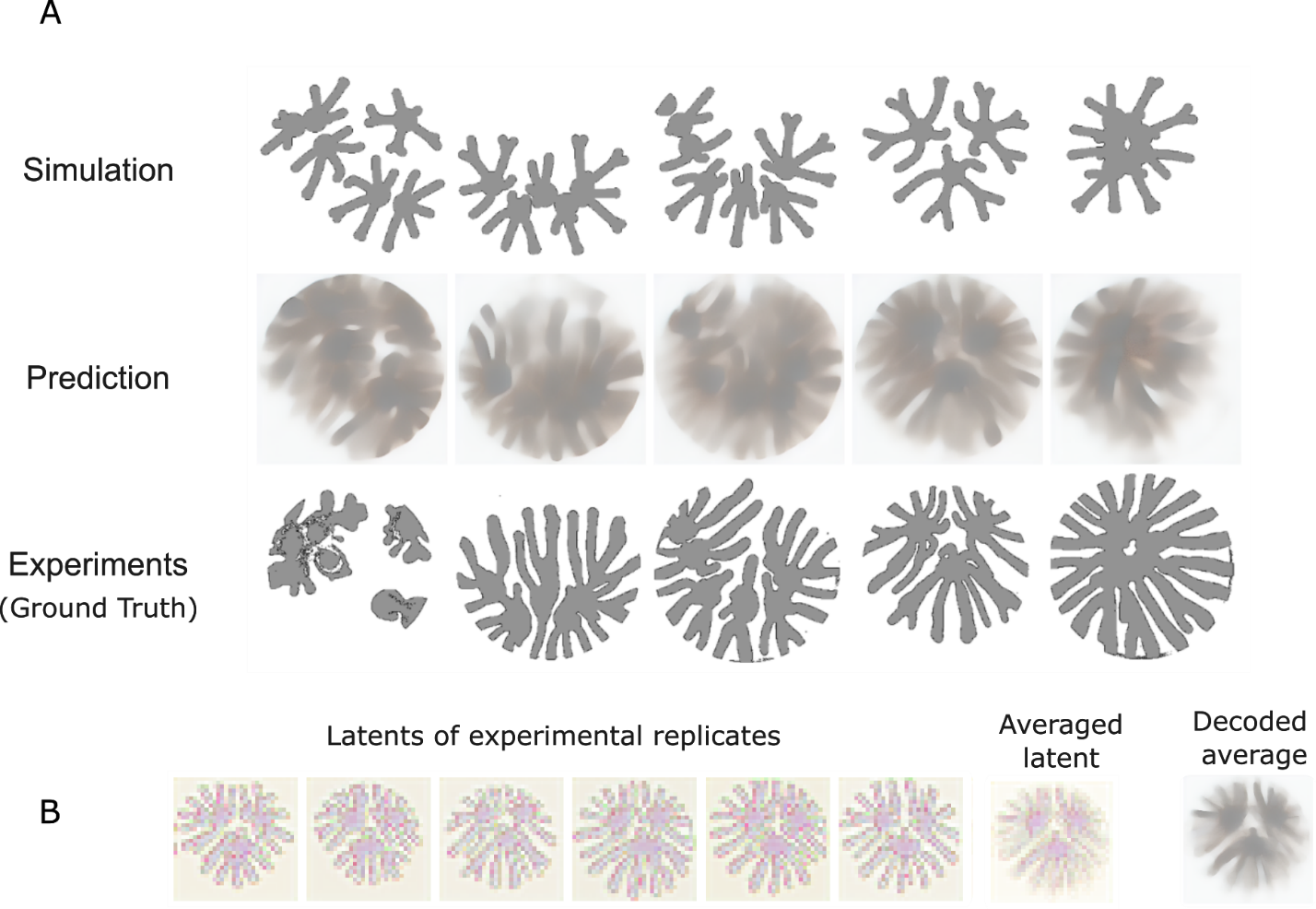

**Supplementary Figure 12***:* **Blurry prediction of experimental patterns from simulations is similar to the averaged image**

(A) *Model performance*: Using simulated patterns as a model input to the ResNet (first row), several different seeding configurations(columns) were tested for prediction of experimental patterns (last row). The model struggles in predicting fine local details and produces a blurry image (middle row).
(B) Latent representations corresponding to 6 experimental replicates of a defined seeding configuration are visualized here using the RGBA scheme (first five columns), along with pixel-wise average latent image (second to last column). Decoding the latents using the SD VAE decoder resulted in a blurry output (last column), similar to the prediction using ResNet shown in (A)

**Supplementary Figure 13**: **Samples of experimental patterns including replicates**

This figure shows a random sample of 100 experimental *Pseudomonas aeruginosa* branching patterns that have been processed using the algorithm described in Supplementary figure 11. We include technical and sometimes biological replicates of the same input seed. The majority of the patterns have 2 technical replicates, while some of the seeding conditions can have up to 6 technical/ biological replicates. Latent compression of these outputs will serve as output or ground truth for the ControlNet diffusion model shown in Figure 5. These images have dimensions of 256 x 256 x 3.

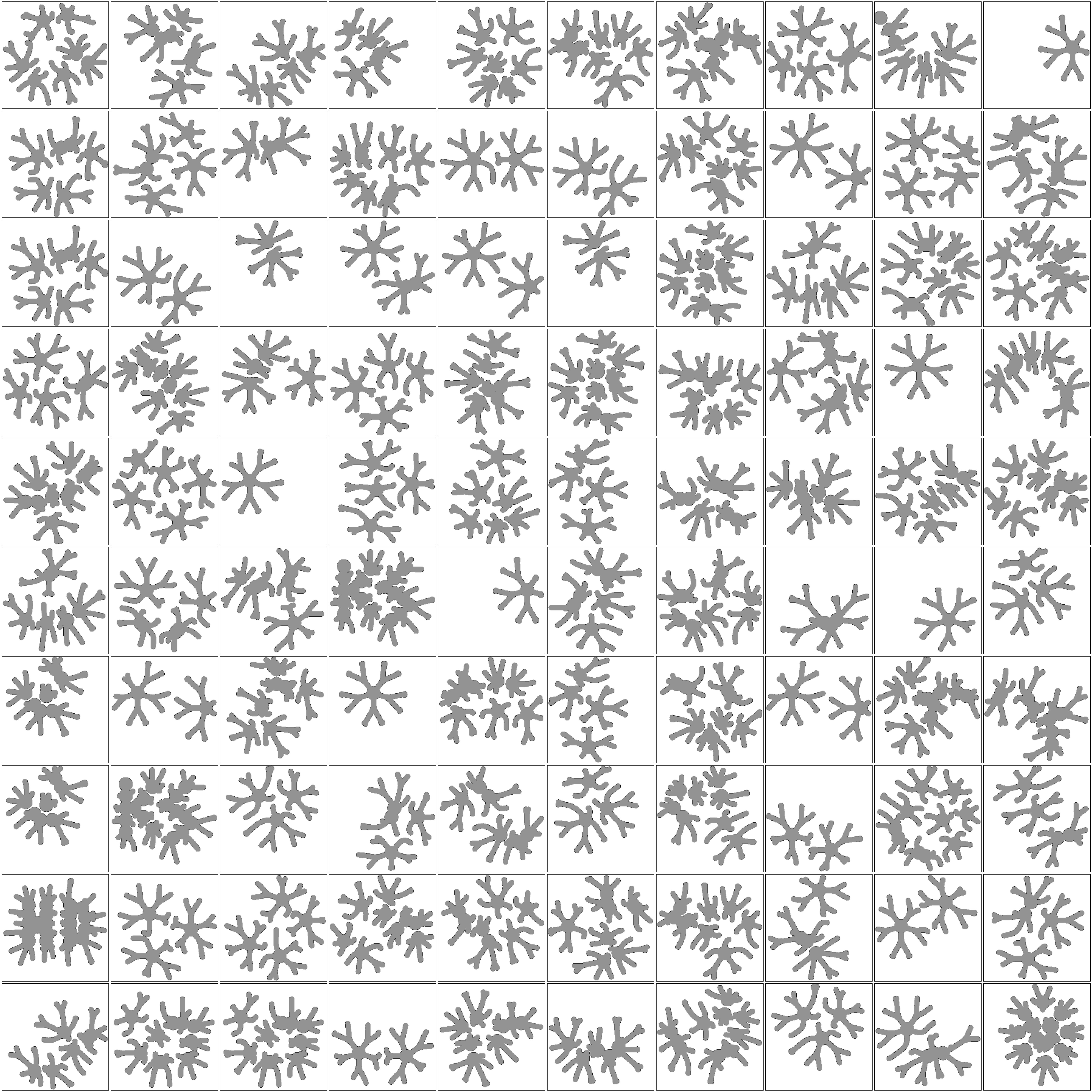

**Supplementary Figure 14**: **Samples of simulated patterns including replicates**

These simulated patterns correspond to the experimental patterns in Supplementary Figure 13. As the simulation branching patterns are deterministic in nature, corresponding to the technical and biological replicates, we include repetitions of the same simulation. Latent compression of these patterns will serve as input or spatial conditioning for the ControlNet diffusion model shown in Figure 5. These images have dimensions of 256 x 256, which are then triplicated to 256 x 256 x 3 before computing the latents.

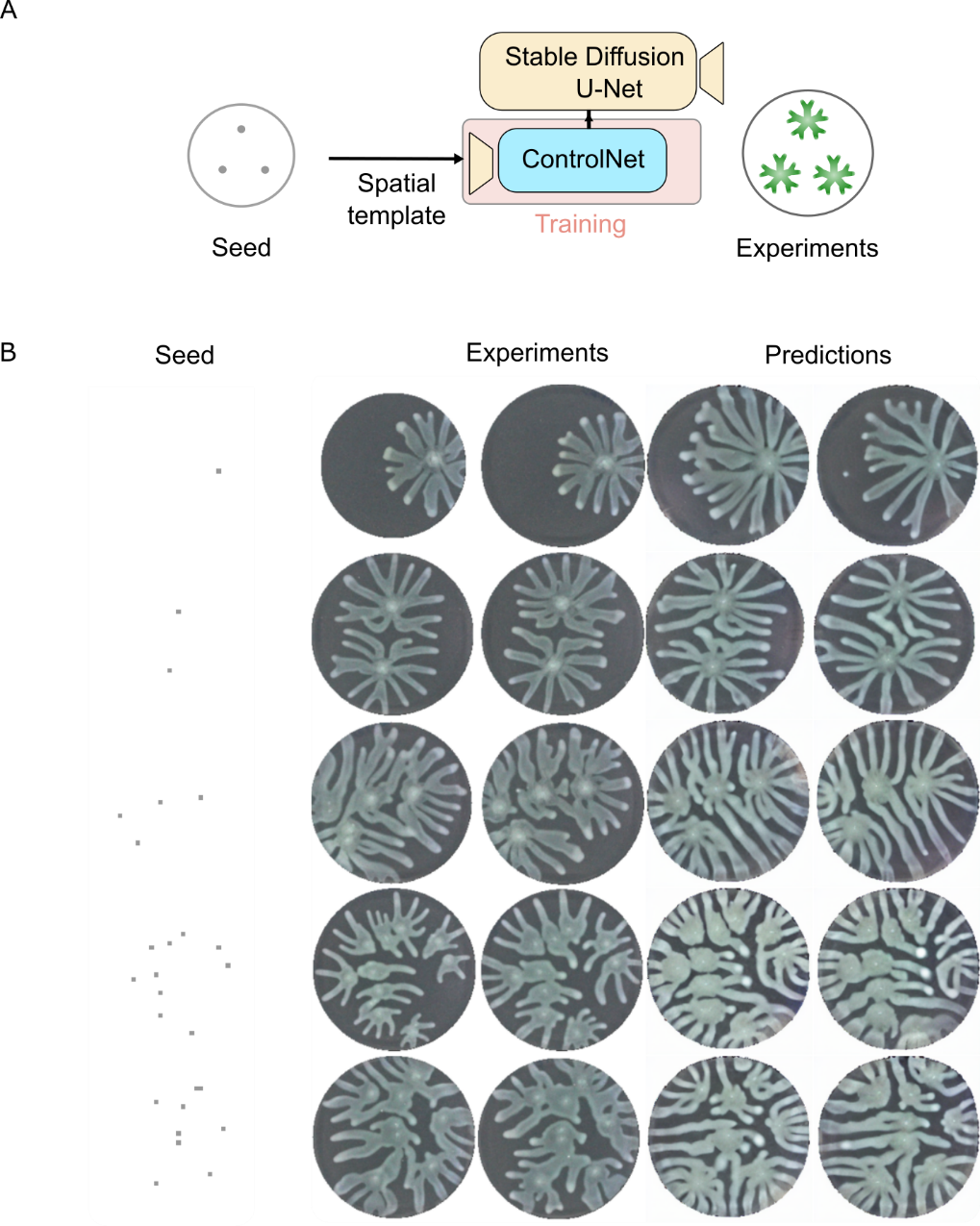

**Supplementary Figure 15: Conditioning on initial seeding configurations as a baseline produced plausible but less simulation-grounded patterns**

A similar pipeline was used as shown in Figure 5, except instead of conditioning on the initial simulations, we conditioned the network on the initial seeding configurations. The training and inference hyperparameters are the same as shown in Figure 5. The quality of prediction is worse, for instance merging of branches (last and second last row) in dense seeding configurations is nonrealistic, with much larger branches and non-consistent branching dynamics compared to the experiments.

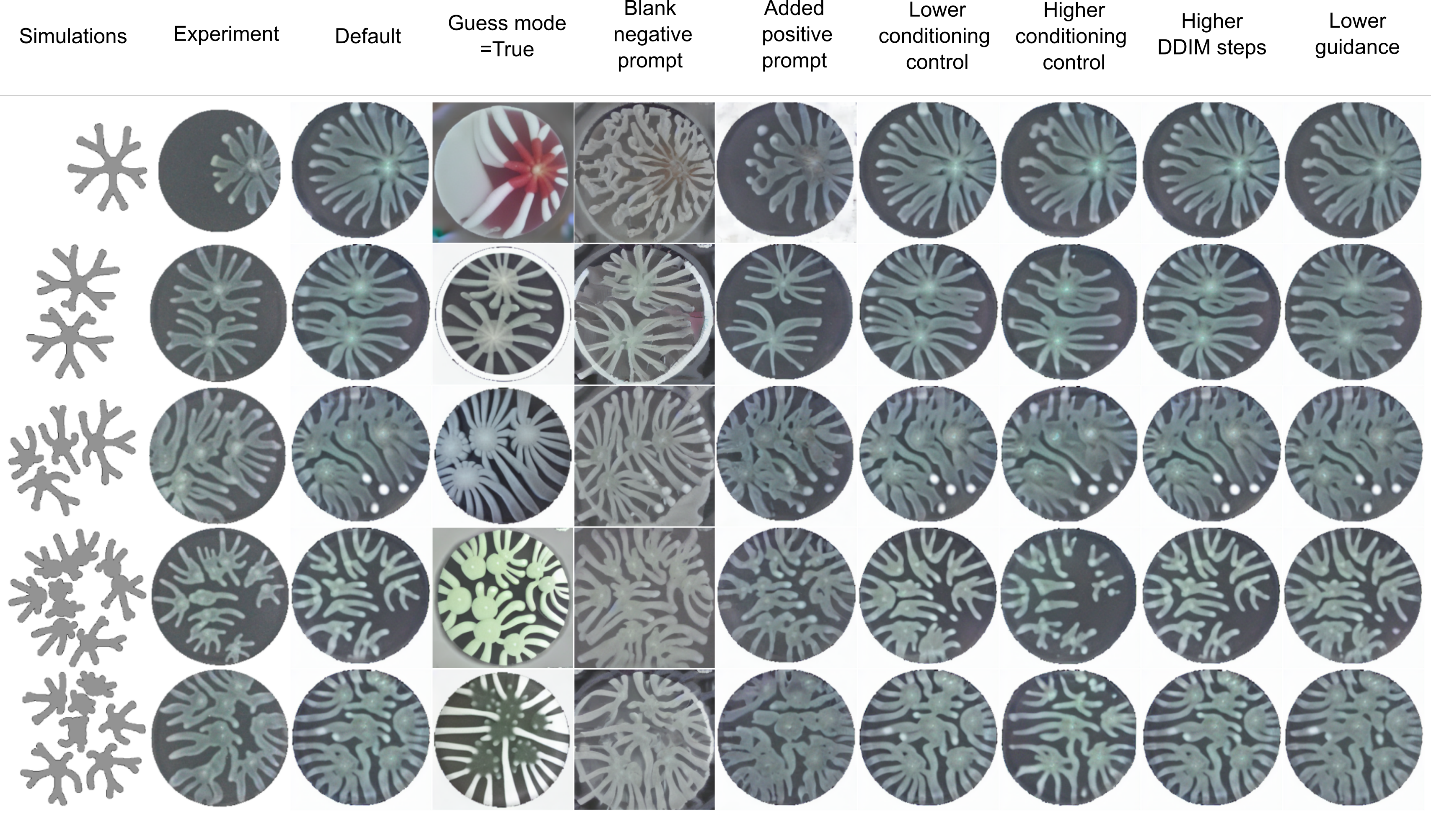

**Supplementary Figure 16**: **Ablation study on controllable parameters in model inference**

During inference, all parameters were kept fixed at the values used in Figure 5 except for the parameter indicated in each column heading (outlined in **Methods**). The first columns show simulated patterns, with different rows indicating patterns from different spatial seeding configurations. These simulations serve as input to the ControlNet model. The second column shows the corresponding experimental patterning pairs (1 replicate). The third column indicates the predictions from the ControlNet model that was used in Figure 5 to provide a baseline estimate. Column 4 shows patterning where the Guess mode parameter is set as True. Overall patterning dynamics look more rigidly structured than experimental patterns, indicating that the conditioning is stronger than needed. Column 5 indicates patterns without the negative prompt used in Figure 5. Removing the negative prompts resulted in worse quality of predictions compared to the baseline. The result of adding positive prompts, shown in Column 6, default ones listed in the ControlNet inference codebase, was also a considerable decrease in experimental plausibility by visual inspection. Columns 7 and 8 show the effect of adding a lower and higher conditional control strength respectively, compared to the baseline. Although the level of changes was not significant compared to the earlier parameters, patterns still looked visibly different, for example more whitish-like simulations in lower strength and distorted plate boundaries and smaller branching area in higher strength. Column 9 indicates a higher number of DDIM diffusion sampling steps, which do not lead to any visible improvement in performance. Column 10 shows a lower guidance value, used in the training of the network. Slight performance decline is visible compared to the baseline, including small singular colony spots away from the patterns (fourth row).

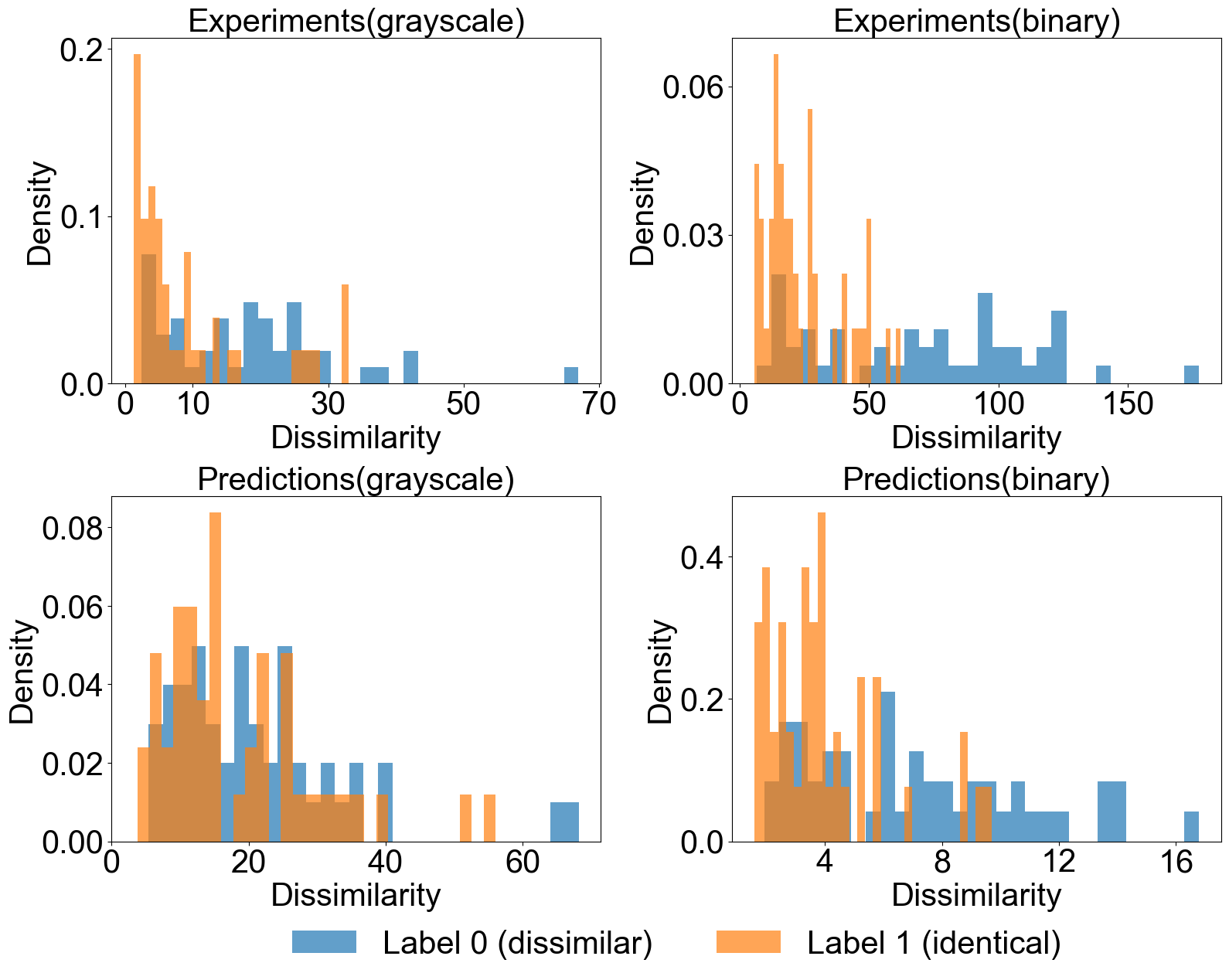

**Supplementary Figure 17**: **Intra condition variation is less than inter condition variation in both experiments and diffusion model predictions.**

A Siamese network was trained on simulated branching patterns to distinguish between identical/non identical patterning pairs. The network was then tested on experiments and diffusion model predictions. Label 0 indicates images from different seeding configurations while Label 1 indicates replicate images. Across both experimental and prediction images, either grayscale or binary thresholded, inter-condition variation, i.e. among non-replicates was higher than intra- condition variation, i.e., among replicates. This indicates although the patterning experiments are stochastic, there is enough information to distinguish between replicate and non-replicate populations.

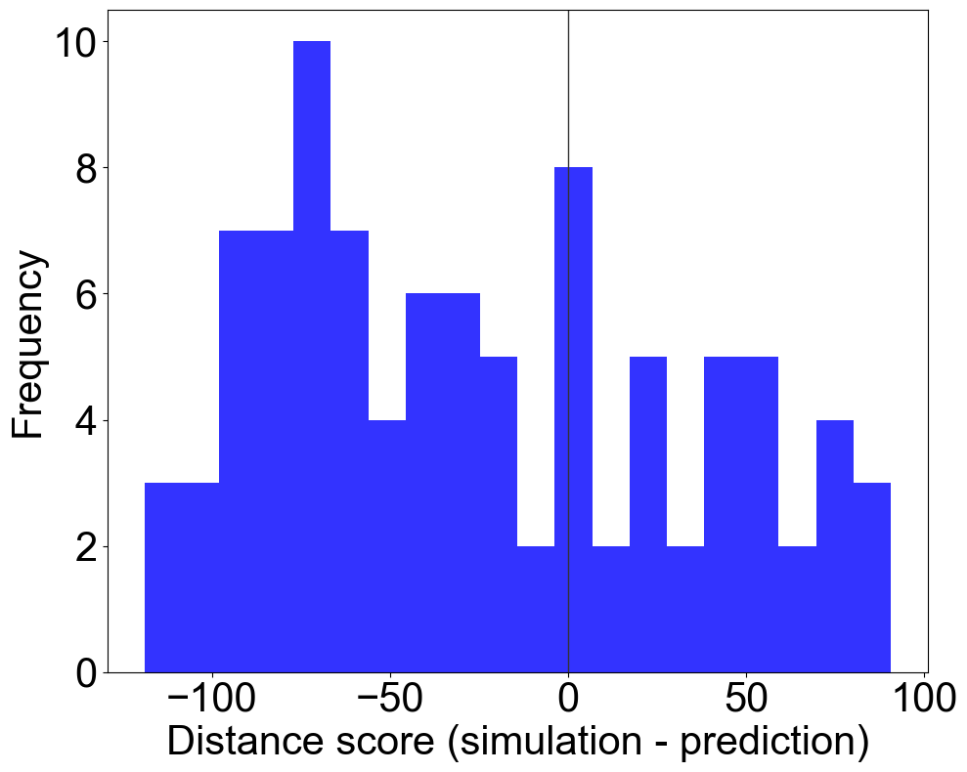

**Supplementary Figure 18**: **Dissimilarity score differences between simulation-experiment and prediction-experiment patterning pairs.**

A Siamese network was trained on simulated branching patterns to distinguish between identical-non identical patterning pairs. The cosine dissimilarity was then calculated to compute the distance between simulation- experiment pattern pairs and compared with prediction-experiment patterning pairs, with the distance score denoting the difference. All images were binary thresholded to test the structural aspect of prediction and remove the confounding factor of color information. For about 1/3^rd^ of the image samples, the prediction samples were closer than simulations in terms of the distance from corresponding experimental samples.

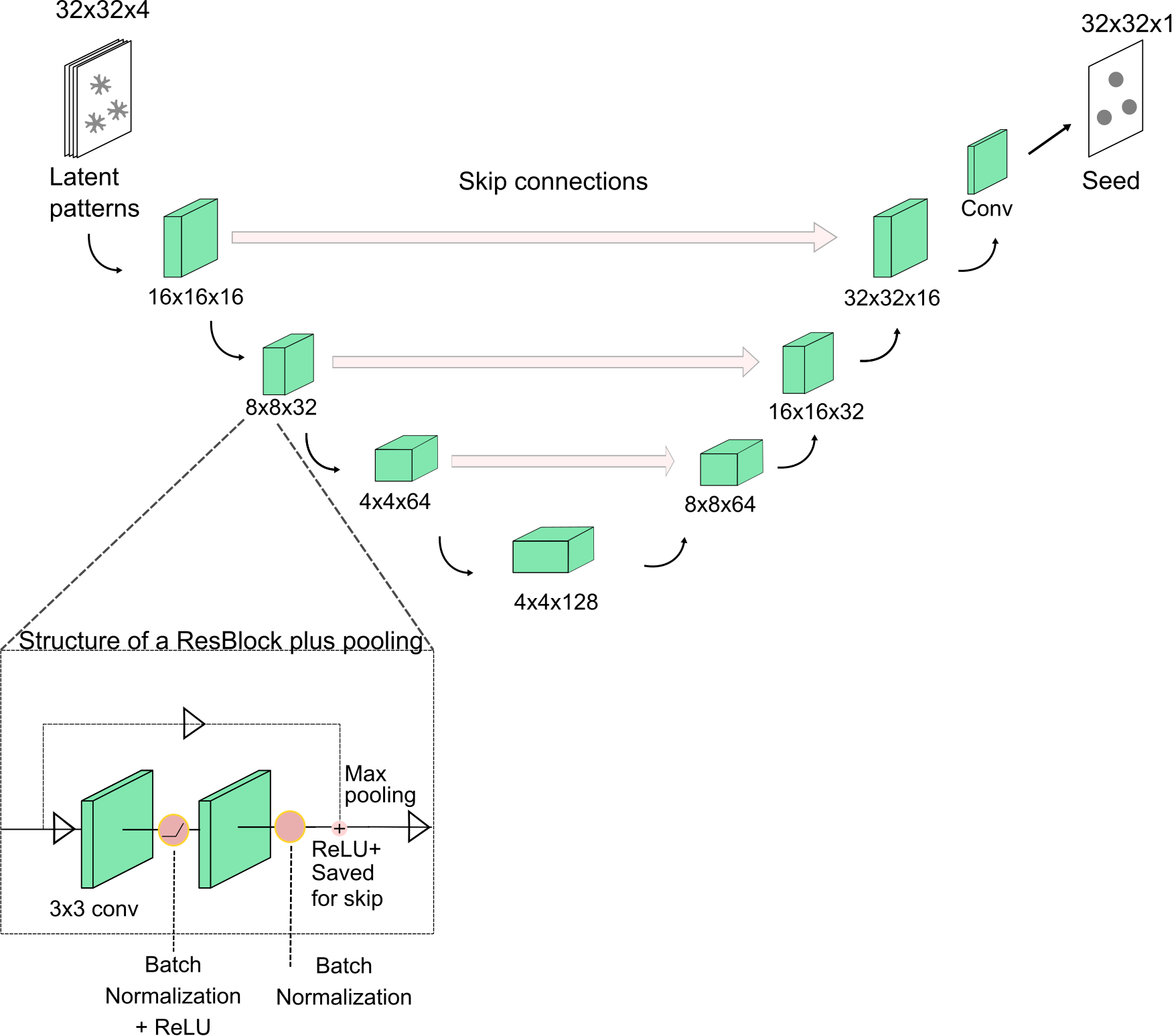

**Supplementary Figure 19:** **U-Net architecture**

The high dimensional input image (256 x 256 x 3) is first compressed using Stable Diffusion VAE Encoder into a latent dimensional representation of dimensions 32 x 32 x 4 for efficient training. The U-Net was then trained to map between compressed experimental/simulated patterns(32 x 32 x 4) and the corresponding seeding configurations(32 x 32 x 1), with contraction path containing three blocks of sizes 16,32 and 64, a bottleneck size of 128, and expansion path containing three blocks of sizes from 64,32,16 before a final convolutional layer is used to decrease the channels to the target image. Each block is composed of a ResBlock followed by a 2x2 max pooling (inset), with the ResBlock having two 3x3 convolutional layers with batch normalization and ReLU. Batch normalization is also applied after the second convolutional layer, and after adding the identity and applying ReLU, outputs are saved for skip connections (before pooling) in the U-Net. Dropout is also implemented between the convolutional layers for regularization. The skip connections are implemented in line with standard U-Net architecture, meant to capture both global and local spatial information. The final output of the network is in logits, and the network is trained with BCE loss for logits plus a weighted Dice term.

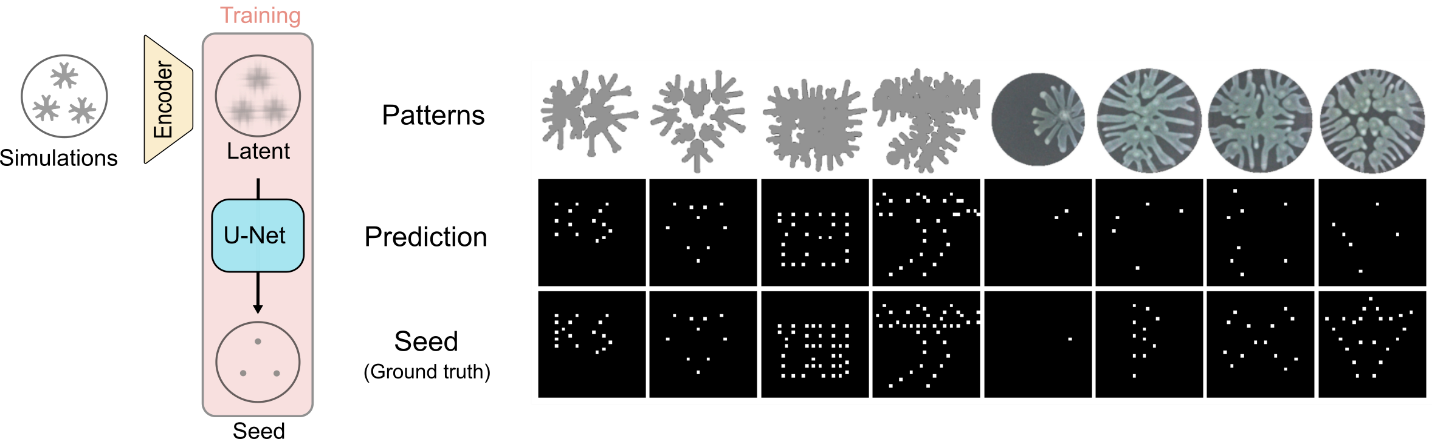

**Supplementary Figure 20: Model trained on simulations fails to generalize to experimental patterns**

The inverse inference pipeline illustrated in Figure 6 is used to infer the initial conditions from the simulated patterns. The training process involves using 90,000 simulated patterns (30,000 patterns x 3 rotational augmentations) which are compressed to latent representations and then a U-Net is used to map from latent representations to the initial seeding configurations. The trained network is then used to infer initial seeds from simulated patterns on the test set (right, first four columns). The model performs well at inferring the initial seeds (Prediction, 2^nd^ row and Ground truth, last row) from the simulated patterns (1^st^ row), even performing well on out of distribution samples- either greater number of seeds or denser seeds (columns 3 and 4). But when the trained model is used on the experimental test dataset, the prediction performance is qualitatively quite worse, both in terms of false positives and false negatives (columns 5 through 8, second row images compared with last row images).

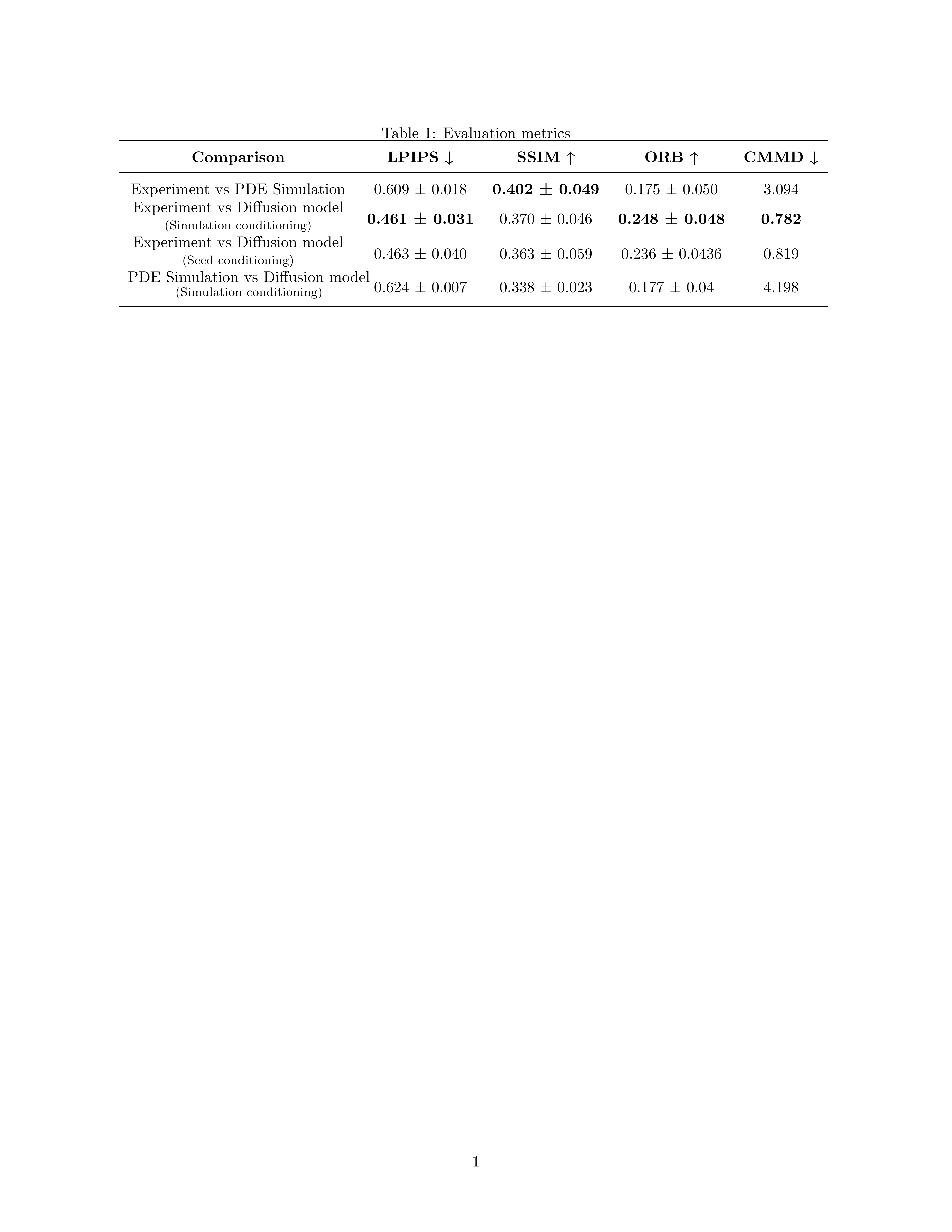

**Supplementary Table 1**: **Evaluation metrics (with mean and standard deviation) on the performance of ControlNet from Figure 5 on the test set.**

Patterns corresponding to seeding configurations used in the test set were compared across simulations, experiments and predictions from the ControlNet model, along with the predictions using the seed conditioning as a baseline. Across most metrics, experiments were found to be closer to the ControlNet predictions conditioned on the simulations compared with simulations. Also, since CMMD does not require paired datasets, we computed the score on experimental images across training vs the test set and found that the score is 0.141.

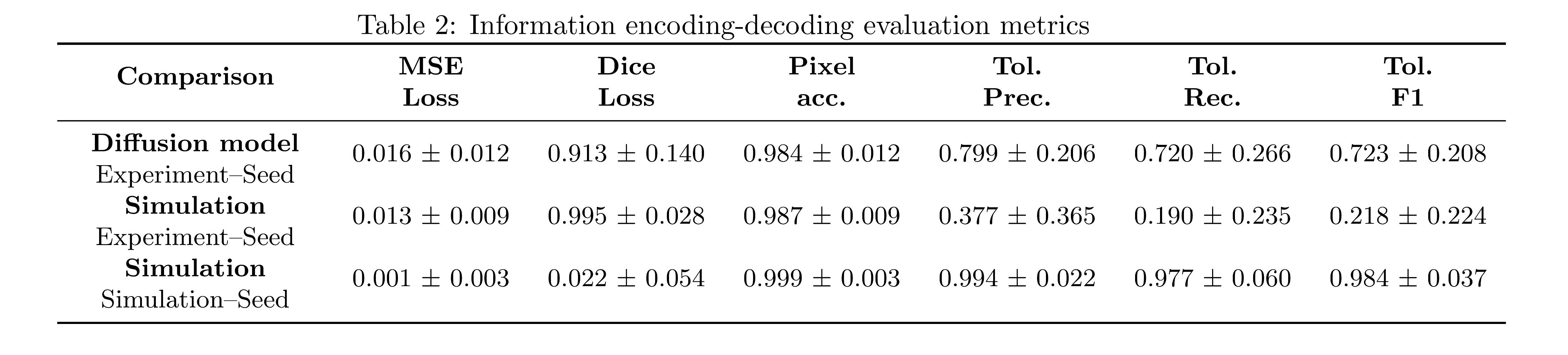

**Supplementary Table 2: Evaluation metrics of model performance for information encoding-decoding task of inferring initial seeds from final patterns**

The performance of different models was tested first on inferring the initial seeds from experimental patterns. Different metrics were computed on the test set, including MSE loss, Dice Loss, Pixel accuracy, Tolerance Precision, Tolerance Recall and Tolerance F1 (tolerance was used to match the points within 4-pixel distance using Hungarian algorithm). Model that is trained on synthetic data using the ControlNet pipeline (first row) has much higher tolerance precision, recall and F1 scores, lower Dice losses, and similar MSE loss and Pixel accuracy when compared with a model that is trained on simulation dataset. As a baseline, the model that is trained on simulations performs much better when the performance is tested on the simulation test set, achieving high values of pixel accuracy, tolerance precision, recall and F1 scores and low values of MSE and Dice losses.
